# Point mutations in the catalytic domain disrupt cellulose synthase (CESA6) vesicle trafficking and protein dynamics

**DOI:** 10.1101/2022.04.04.487015

**Authors:** Lei Huang, Weiwei Zhang, Xiaohui Li, Christopher J. Staiger, Chunhua Zhang

## Abstract

Cellulose, as the main component of the plant cell wall, is synthesized by a multimeric protein complex named the cellulose synthase (CESA) complex or CSC. In plant cells, CSCs are transported through the endomembrane system to the PM, but how catalytic activity or conserved motifs around the catalytic core domain influence vesicle trafficking or protein dynamics is not well understood. Here, we used a functional YFP-tagged AtCESA6 and site-directed mutagenesis to create 18 single amino acid replacement mutants in key motifs of the catalytic domain including DDG, DXD, TED and QXXRW, to comprehensively analyze how catalytic activity affects plant growth, cellulose biosynthesis, complex formation, as well as CSC dynamics and trafficking. Plant growth and cellulose content were reduced by nearly all mutations. Moreover, mutations in most conserved motifs reduced the speed of CSC movement in the PM as well as delivery of CSCs to the PM. Interestingly, the abundance of YFP-CESA6 in the Golgi apparatus was increased or reduced by mutations in DDG and QXXRW motifs, respectively. Post-Golgi trafficking of CSCs was also differentially perturbed by these mutations and, based on these phenotypes, the 18 mutations could be divided into two major groups. Group I comprises mutations causing significantly increased fluorescence intensity of YFP-CESA6 in Golgi with either an increase or no change in the abundance of cortical small CESA-containing compartments (SmaCCs). In contrast, Group II represents mutations with significantly decreased fluorescence intensity of YFP-CESA6 in Golgi and/or reduced SmaCC density. In addition, two Group II mutations in the QXXRW motif reduced CSC assembly in the Golgi. We propose that Group I mutations cause CSC trafficking defects whereas Group II mutations, especially in the QXXRW motif, affect normal CSC rosette formation in the ER or Golgi and hence interfere with subsequent CSC trafficking. Collectively, our results demonstrate that the catalytic domain of CESA6 is essential not only for cellulose biosynthesis, but also CESA complex formation, protein folding and dynamics, vesicle trafficking, or all of the above.

**One sentence summary:** A comprehensive mutational analysis of the catalytic domain of Arabidopsis CESA6 reveals distinct roles for conserved motifs in CSC vesicle trafficking, protein complex formation, or protein dynamics

## Introduction

The plant cell wall provides a tough but flexible barrier to the outside environment and plays important roles in plant growth and development, as well as protecting plants from biotic and abiotic stresses (McFarlane et al., 2014; Zhang et al., 2021a). The cell wall consists of a network of polysaccharides including cellulose, hemicellulose and pectin. Cellulose is the main polysaccharide of the primary cell wall and comprises linear β-1,4-linked glucose polymers arrayed into semi-crystalline microfibrils that act as the primary load-bearing component of the cell wall. As an important and abundant renewable natural resource, cellulose is produced predominantly by plants, algae and certain bacteria, and makes major contributions to food, fiber and biofuel industries (Somerville, 2006).

Cellulose is synthesized at the plasma membrane (PM) by integral membrane protein complexes called cellulose synthase (CESA) complexes or CSCs. These complexes were first visualized as 6-fold symmetry “rosette” structures in freeze fracture samples by electron microscopy (Mueller and Brown, 1980; Haigler and Brown, 1986). The Arabidopsis (*Arabidopsis thaliana*) genome has 10 *CESA* genes and, based on genetic and biochemical assays, each CSC contains three different CESA isoforms with a 1:1:1 molecular ratio (Desprez et al., 2007; Persson et al., 2007). Depending on the type of cell wall being produced, CESAs are further divided into primary and secondary cell wall complexes. The primary cell wall CSCs in Arabidopsis contain CESA1, CESA3 and CESA6 or CESA6-like proteins, whereas the secondary cell wall CSCs are CESA4, CESA7 and CESA8 (Desprez et al., 2007; Persson et al., 2007). The cryo-electron microscopy structure of *Populus tremula x tremuloides* (*Ptt*) CESA8 suggests that each CSC rosette is likely to comprise six trimers of CESA and therefore 18 CESAs in total (Purushotham et al., 2020). Each CESA protein in a CSC is thought to produce a single glucan chain and many of these are linked by hydrogen bonds to form cellulose microfibrils.

All CESAs are integral membrane proteins and share an overall domain architecture: an N-terminal cytoplasmic domain, seven transmembrane spans, and a glycosyltransferase type-2 (GT2) domain located between the 2^nd^ and 3^rd^ transmembrane domains (Purushotham et al., 2020; Zhang et al., 2021c). The GT2 domain contains the catalytic core of CESA and is responsible for synthesizing β-1,4-glucans using UDP-glucose as the substrate. Several important motifs are located in the catalytic domain including the plant-conserved region (PCR), the class-specific region (CSR), as well as canonical motifs such as DDG, DXD, TED and QXXRW motifs (Vergara and Carpita, 2001; Purushotham et al., 2020; Daras et al., 2021). The PCR domain creates an electropositive trimer interface through hydrophobic contacts with neighbors and facilitates CESA trimerization (Purushotham et al., 2020). The cysteine-rich CSR domain is thought to facilitate CESA membrane trafficking via S-acylation (Kumar et al., 2016). The DDG, DXD, TED and QXXRW motifs in the catalytic domain are conserved among plant and bacteria and they participate in donor and acceptor UDP-glucose binding, glucan chain polymerization and elongation (Morgan et al., 2013; Morgan et al., 2016; Purushotham et al., 2020). A recent crystal structure of the AtCESA3 catalytic domain reveals the ability to form homodimers through contacts between β-strand 6 (β6) and α-helix 9 (α9) of adjacent subunits and bimolecular fluorescence complementation suggests that homodimers form in the Golgi apparatus (Qiao et al., 2021). This could represent a catalytically-inactive structure of CESA subunits and perhaps contributes to CSC assembly, as proposed previously (Atanassov et al., 2009).

Originally, the subcellular distribution of CSC rosettes at the PM, Golgi cisternae and Golgi-derived vesicles was documented by freeze-fracture electron microscopy of *Zinnia* mesophyll cells (Haigler and Brown, 1986). The advent of fluorescent protein tags and high resolution confocal microscopy of living plant cells, along with a functional YFP-tagged Arabidopsis CESA6, confirmed and extended the subcellular location of CESA and CSCs to include descriptions of their dynamic trafficking and behavior at the PM (Paredez et al., 2006). Functional YFP-AtCESA6 is present in the Golgi apparatus, in vesicles some of which are named small CESA-containing compartments (SmaCCs) or microtubule-associated cellulose synthase compartments (MASCs), as well as in the PM (Paredez et al., 2006; Crowell et al., 2009; Gutierrez et al., 2009). As a typical secreted protein, the current paradigm holds that CESA proteins are synthesized in the endoplasmic reticulum (ER), transported to the Golgi for CSC complex assembly, trafficked to the trans-Golgi network (TGN) and packaged into post-Golgi vesicles for delivery to the PM (Polko and Kieber, 2019; Blichfeldt et al., 2022). The cortical SmaCCs are thought to be involved in both the secretion and internalization of CSCs, although their exact identity remains somewhat elusive (Crowell et al., 2009; Gutierrez et al., 2009). In recent years, through CESA co-expression analysis, multiple CESA-associated proteins have been identified and shown to participate in CESA trafficking (Persson et al., 2005; Sanchez-Rodriguez et al., 2012; Endler et al., 2015; Zhang et al., 2016; McFarlane et al., 2021; Vellosillo et al., 2021; Gu and Rasmussen, 2022). CSC exocytosis is regulated by multiple components and partners, such as the exocyst complex (Zhu et al., 2018), actin and myosin XI (Sampathkumar et al., 2013; Zhang et al., 2019; Zhang et al., 2021b), Rab GTPases (RabH1b; He et al., 2018), as well as other novel players. For example, SHOU4 interacts with CESAs and negatively regulates the exocytosis of CSCs to the PM (Polko et al., 2018); the plant-specific protein PATROL1 assists with vesicle tethering or fusion to the PM (Zhu et al., 2018); and TRANVIA is associated with CSCs in the secretory pathway and facilitates delivery to the PM (Vellosillo et al., 2021). Interestingly, two different CESA-interacting proteins, STELLO1/2 and 7TM localize primarily to the Golgi or TGN and play a role in cellulose biosynthesis and abundance of CSCs at the PM (Zhang et al., 2016; McFarlane et al., 2021). At the end of their life or during recycling, PM-localized CSCs are internalized through clathrin-mediated endocytosis to TGNs/early endosomes (EEs), which maintains CSC homeostasis at the PM (Bashline et al., 2013; Bashline et al., 2015; Sanchez-Rodriguez et al., 2018; Zhu and McFarlane, 2022).

Cellulose biosynthesis inhibitors (CBIs) are widely used small molecules that have greatly enhanced our understanding of the mechanism of cellulose synthesis in plant cells (Larson and McFarlane, 2021). Recently, we identified and characterized a novel CBI named Endosidin 20 or ES20, which targets the catalytic site of CESA6 and inhibits cellulose synthesis, CSC motility at the PM, as well as CSC trafficking (Huang et al., 2020), suggesting that the catalytic domain not only contributes to enzymatic activity but also affects efficient trafficking of CESA and protein dynamics at the PM. Similarly, AtCESA1^D605N^ (*any1*) with a point mutation in the catalytic domain showed reduced CSC motility at the PM (Fujita et al., 2013) and AtCESA6^D395N^, mutated in the DDG motif, showed markedly reduced fluorescence of CSCs at the PM and Golgi, and was described to localize in the ER (Park et al., 2019). Surprisingly, little is known about how the conserved catalytic core domain of CESA contributes to CSC trafficking, delivery to the PM, or protein dynamics.

To further elucidate the role of the catalytic domain in enzymatic activity, CSC trafficking and membrane dynamics, a comprehensive set of 18 single amino acid replacement mutations were created in YFP-CESA6 by site-directed mutagenesis and transformed into the *CESA6* null mutant *prc1-1*. Using high spatiotemporal resolution microscopy and quantitative image analysis, we found that multiple mutations in the DXD, TED and QXXRW motifs, but not the DDG motif, caused reduced motility of PM-localized CSCs. Further, 16 of 18 mutations affected CSC trafficking to the PM and exhibited reduced CSC delivery rate to the PM as well as deficiencies in at least one of the Golgi-to-PM trafficking steps. Pairwise correlation analyses indicated that these trafficking phenotypes were mainly associated with defects first visible in the Golgi but exhibited two distinct phenotypes implying different mechanisms. Collectively, our results demonstrate that the CESA6 catalytic domain is not only indispensable for enzymatic activity, but also likely contains information for trafficking regulation and/or functional CSC formation in the ER or Golgi.

## Results

### Mutagenesis of the AtCESA6 catalytic domain reveals key motifs that contribute to hypocotyl growth and cellulose biosynthesis

Previously, we identified 15 AtCESA6 mutants that are insensitive to ES20 treatment and named them *es20r* mutants (Huang et al., 2020). In a subsequent screen, another *es20r* mutant was identified with a leucine to phenylalanine replacement at amino acid 1029 (L1029F). In total, 16 ES20-insensitive AtCESA6 mutants were identified by forward genetic screens of EMS-mutagenized populations (Supplemental Figure S1; Supplemental Table S1). Through homology modeling, molecular docking and biochemical assays, we demonstrated that the predicted ES20-binding pocket on AtCESA6 overlaps with key motifs in the catalytic domain, such as DDG, DXD, TED and QXXRW, and the majority of *es20r/cesa6* mutants were mutated in or near these motifs (Huang et al., 2020; Figure 1, A and B). Our live-cell imaging and quantitative cell biology studies revealed that ES20 treatment affects the motility of CSCs in the plane of the PM, probably by interfering with the catalytic activity of CESAs; in addition, ES20 treatment reduces the trafficking of CSCs to the PM (Huang et al., 2020). These results suggest that the ES20-targeted motifs in CESA6 may play important roles in both CESA catalytic activity and CSC trafficking through the endomembrane system.

**Figure 1.**
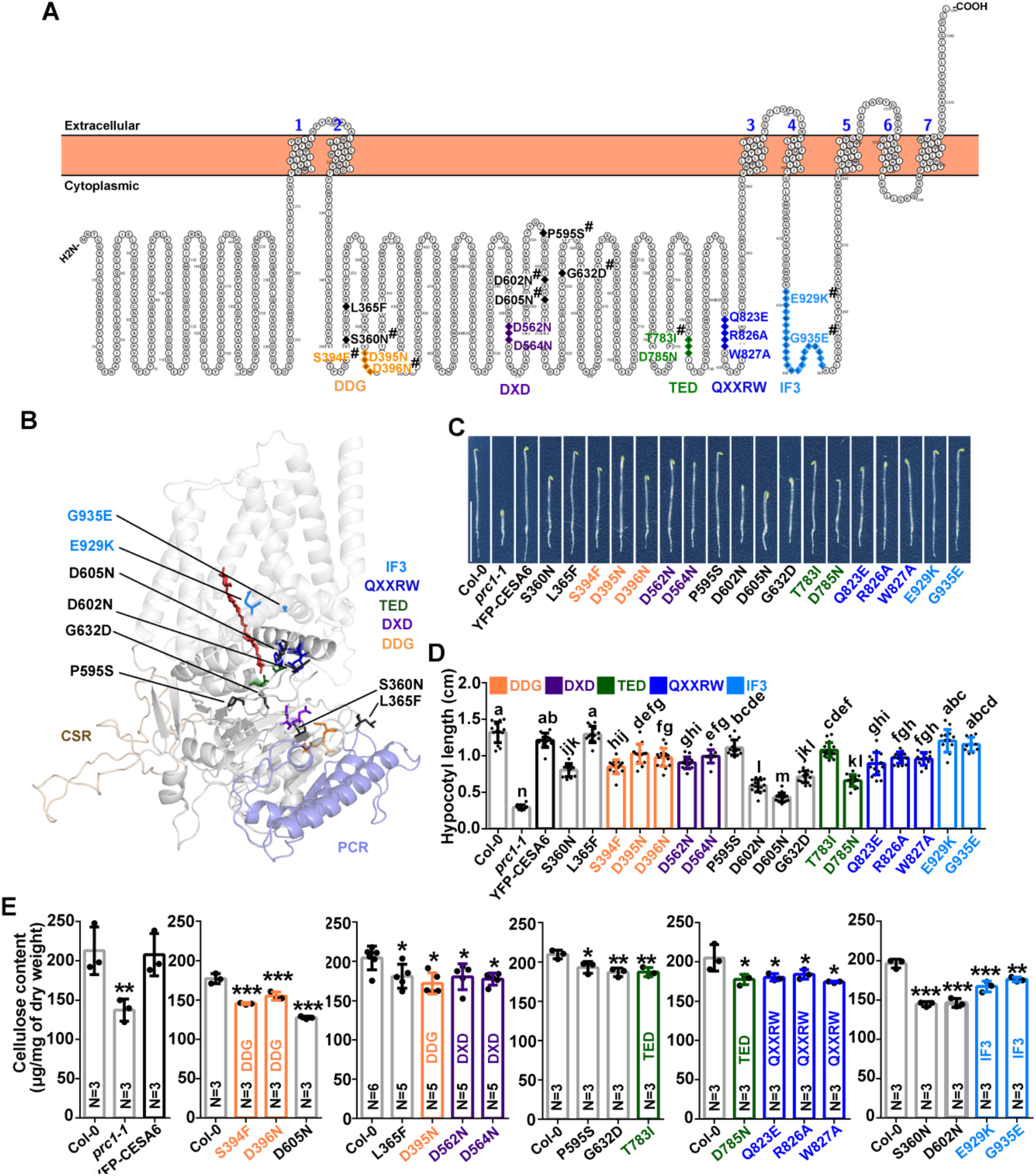
Mutations in key motifs of the CESA6 catalytic domain affect hypocotyl growth and cellulose biosynthesis. **(A)** Topology of AtCESA6. The position of CESA6 amino acid changes introduced by site- directed mutagenesis are indicated. Mutated amino acids in the conserved catalytic core DDG, DXD, TED, QXXRW as well as Interfacial helix 3 (IF3) motifs are highlighted with colors defined in the legend and used throughout the subsequent figures. Mutations outside of these five motifs, but that recapitulate mutations found in the original set of *es20r* mutants (Huang et al., 2020), are shown in dark grey. # Denotes amino acid change/mutation discovered in *es20r* screen and recreated in YFP-CESA6 complementation line (Huang et al., 2020). **(B)** Homology model of full-length AtCESA6 with mutated amino acids highlighted. Modeling of full-length AtCESA6 was performed using the SWISS-MODEL server using the structure of PttCESA8 (Protein Data Bank ID: 6WLB) as template. An elongating glucan chain is depicted in red. Ribbon diagram of the secondary structure elements and the peptide backbone are shown in light gray. **(C)** Representative 3-d-old dark grown seedlings of transgenic plants expressing wild-type or mutated YFP-CESA6. Scale bar: 1 cm. **(D)** Hypocotyl length measured from 3-d- old dark grown seedlings as described in C. Data represent mean ± SD (n ≥ 11 seedlings per genotype). Statistically significant differences were determined using one-way analysis of variance (ANOVA) followed by Tukey’s multiple comparisons test. Different letters indicate significant differences between groups (p < 0.05). **(E)** Crystalline cellulose content in hypocotyl cell walls prepared from 7-d-old dark grown plants expressing YFP-CESA6 with mutations in key motifs. Cell walls prepared from wild-type (Col-0) seedlings, YFP-CESA6 expressed in *prc1-1* complementation line, and homozygous *prc1-1/cesa6* mutant serve as controls. Mutations in the DDG, DXD, TED, QXXRW and IF3 motifs are highlighted with corresponding colors. Data represent mean ± SD. Dots on each bar represent a biological replicate prepared from an independent batch of seedlings. Each biological replicate value was obtained by calculating the mean value of at least three technical replicates. * indicates p < 0.05, ** indicates p < 0.01, and *** indicates p < 0.001 by one-way ANOVA followed by Dunnett’s multiple comparisons test compared with wild type.

To test this hypothesis and better understand the function of various motifs and key amino acids in the CESA catalytic domain and elsewhere on the overall structure, we generated complementation lines by introducing YFP-tagged AtCESA6 harboring single amino acid replacement mutations into the null mutant *prc1-1/cesa6*. Ten *es20r* mutations in or near the catalytic site of AtCESA6 were chosen to create complementation lines (Huang et al., 2020; Supplemental Table S1; Supplemental Figure S1; Supplemental Figure S2; Figure 1, A and B). In addition, we mutated eight additional amino acids in the conserved catalytic motifs including DDG, DXD, TED and QXXRW and introduced them into *prc1-1* (Supplemental Table S1; Supplemental Figure S2; Figure 1, A and B). Collectively, a total of 18 single amino acid changes were introduced into the functional YFP-AtCESA6 fusion protein by site-directed mutagenesis and all constructs were driven by the *AtCESA6* promoter. We quantified *AtCESA6* transcript levels in wild type and mutants by RT-qPCR and found that *AtCESA6* transcript levels in 13 mutants were comparable to wild type, whereas five mutants showed decreased *AtCESA6* transcript levels including S360N, L365F, D562N, G632D and E929K (Supplemental Figure S3, A and B). In addition, by using semi-quantitative immunoblotting with anti-GFP, we found that the normalized expression levels of YFP-AtCESA6 in the wild-type complementation line as well as the 18 transgenic lines harboring single amino acid changes were comparable (Supplemental Figure S3, C to F), reducing the likelihood that any phenotypes were caused by variations in CESA6 protein levels.

Because a majority of the 18 YFP-CESA6 mutated transgenic lines targeted key motifs of the catalytic domain (Figure 1, A and B), and because CESA mutants typically exhibit visible plant growth phenotypes (Harris et al., 2012; Fujita et al., 2013), we first tested whether the YFP-CESA6 mutants could complement the seedling growth phenotype of *prc1-1*. Here, hypocotyl length under dark growth conditions was evaluated. The wild-type YFP-CESA6 as well as four mutated YFP-CESA6 constructs, including L365F, P595S and interfacial helix 3 (IF3) motif mutations (E929K and G935E), fully rescued the *prc1-1* hypocotyl growth phenotype, whereas the other 14 mutated AtCESA6 complementation lines including mutations in the DDG, DXD, TED and QXXRW motifs only partially rescued the growth phenotype (Figure 1, C and D). We also analyzed the full set of the original untagged AtCESA6 EMS-induced mutants and approximately half of them showed hypocotyl growth defects (Supplemental Figure S1, A and B).

The CESA catalytic domain utilizes UDP-glucose as the substrate to synthesize β-1,4-glucan chains of cellulose. The canonical DDG, DXD, TED and QXXRW motifs are involved in substrate binding, glucan chain elongation and translocation (Morgan et al., 2013; Morgan et al., 2016; Purushotham et al., 2020). Moreover, previous results show that ES20 treatment results in significantly reduced cellulose content in the cell wall (Huang et al., 2020). Therefore, we hypothesized that mutations in the ES20-targeted motifs or key motifs of the catalytic domain would cause reduced cellulose synthesis as summarized in Supplemental Table S2. We analyzed crystalline cellulose content in hypocotyl cell wall fractions prepared from the mutated YFP-CESA6 transgenic lines. As shown in Figure 1E, only the wild-type YFP-CESA6 complementation line completely rescued the cellulose biosynthesis deficiency of *prc1-1*, whereas all of the mutated AtCESA6 complementation lines showed a significant reduction in cellulose content, ranging from 7.9% (P595S) to 27.9% (D605N) compared with wild type. We also quantified the crystalline cellulose content in hypocotyl cell wall fractions from the original *es20r* EMS mutant lines and found that only three mutants had crystalline cellulose content comparable to wild type, namely D602N, S818T and G935E, whereas the remaining 13 mutants exhibited significantly reduced cellulose content (Supplemental Figure S1C). Collectively, these data demonstrate that key motifs in the AtCESA6 catalytic domain contribute to hypocotyl growth and cellulose biosynthesis.

### Linear motility of CSCs in the plane of the PM is reduced by mutations in the DXD, TED and QXXRW motifs

To further explore the mechanisms that underpin how these AtCESA6 mutations affect cellulose synthesis, we tested whether the linear movement or motility of CSCs in the plane of the PM was affected. The motility of CSCs is attributed to the catalytic activity of CESA which polymerizes and extrudes glucan chains to form cellulose microfibrils in the cell wall; therefore, the speed of CSC translocation in the PM is often used as a surrogate for CESA catalytic activity (Paredez et al., 2006). We hypothesized that mutations in ES20-targeted motifs, especially those located in key motifs of the catalytic domain, would affect CESA catalytic activity and thus reduce the motility of CSCs. Etiolated hypocotyl epidermal cells from mutated YFP-CESA6 transgenic lines were imaged with spinning disk confocal microscopy (SDCM) and the speed of CSCs in the PM of actively elongating cells was analyzed using kymographs, as described previously (Zhang et al., 2019; Huang et al., 2020). As shown in Figure 2 and Supplemental Figure S4, 12 out of 18 AtCESA6 mutation lines showed significantly reduced mean speed compared with wild-type YFP-CESA6 in the *prc1-1/cesa6* background, which translocated at ~ 200 nm/min. We analyzed two or three different mutant lines each for the DXD, TED, and QXXRW motifs and all of them showed significantly reduced CSC speed, with reductions ranging from 10 to 50%. In addition, P595S, D602N, D605N and G632D also caused 15 to 50% reductions in CSC speed, although they do not map to any previously described catalytic motif. For the IF3 motif, mutation of G935E but not E929K showed significantly reduced CSC speed. In contrast, none of the three mutations (S394F, D395N and D396N) in the DDG motif showed a reduction of CSC speed.

**Figure 2.**
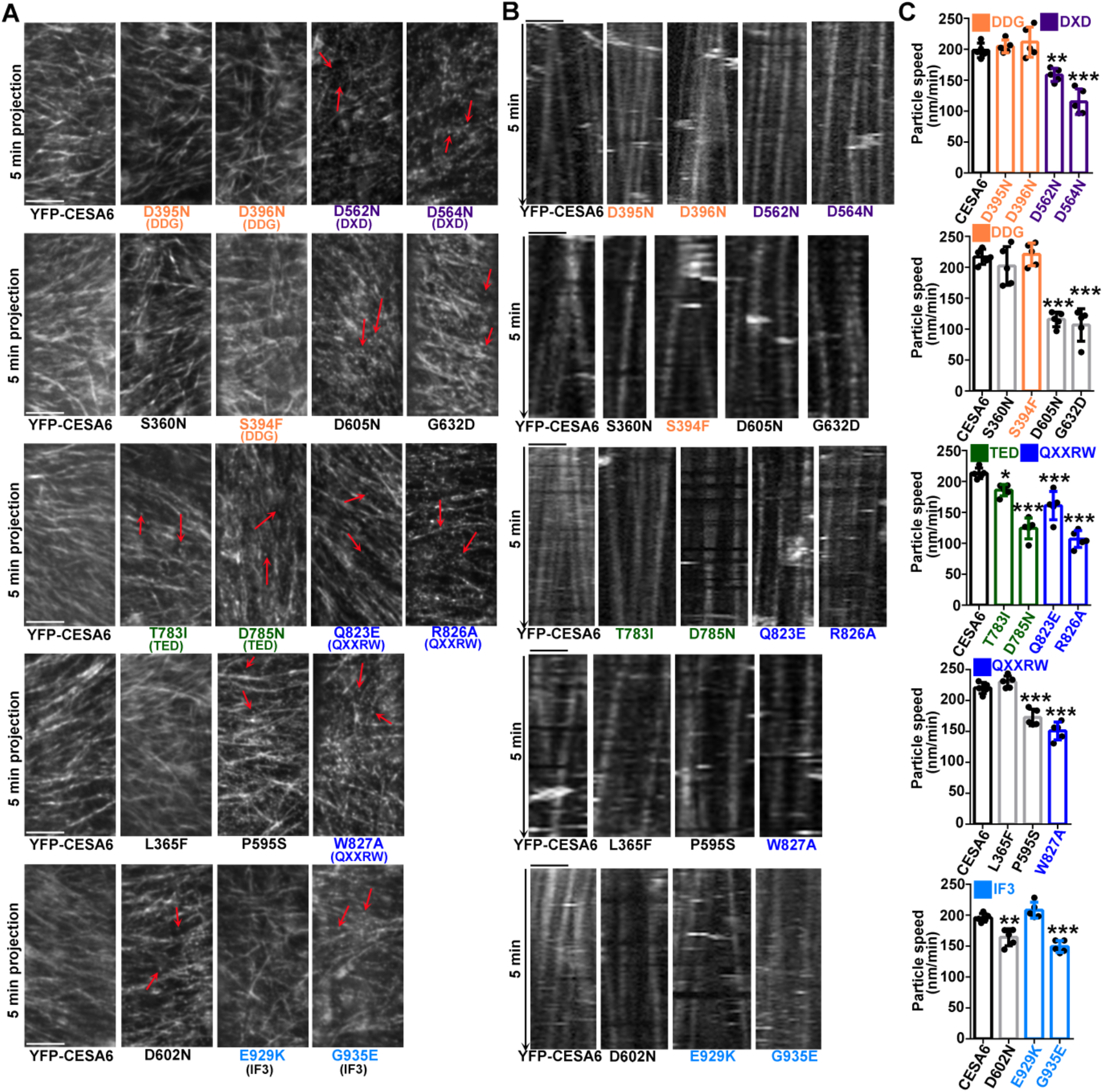
CSC motility is reduced by mutations in the DXD, TED and QXXRW motifs. **(A)** Representative time projections prepared with average intensity images from a time-lapse series of YFP-CESA6 particles at the PM in etiolated hypocotyl epidermal cells of transgenic plants expressing mutated CESA6. Red arrows highlight static CSC particles present in some mutants. Scale bars: 5 μm. **(B)** Representative kymographs of the trajectories of YFP-CESA6 particles from seedlings as described in A. Scale bars: 5 μm. **(C)** Quantification of YFP-CESA6 particle speed. Data represent mean ± SD (n = 5 seedlings per sample, with more than 50 CSC trajectories analyzed for each seedling). * indicates p < 0.05 and *** indicates p < 0.001 by one-way ANOVA followed by Dunnett’s multiple comparisons test compared with wild-type YFP-CESA6 complementation line.

In the 12 mutant lines with reduced CSC speed, we observed a population of CSCs at the PM that were completely static (Figure 2A; red arrows). The insertion of single CSC particles at the PM can be visualized with high spatiotemporal resolution live-cell imaging and the dynamic behavior of CSCs in a typical insertion process follows a specific pattern (Gutierrez et al., 2009; Zhang et al., 2019; Zhang and Staiger, 2022). After a CSC particle arrives at the PM insertion site, it initially pauses for ~ 80 s which is assumed to encompass the tethering, docking, fusion and possibly activation steps. The pause phase is followed by a period of steady movement during which the CSC actively translocates in the PM plane at a relatively constant speed to synthesize cellulose microfibrils (Gutierrez et al., 2009; Zhang et al., 2019; Duncombe et al., 2021; Zhang et al., 2021b; Figure 3, A and B). To confirm that the static CSCs in the CESA6 mutants are non-motile from the beginning of their insertion at the PM, we performed single-particle CSC insertion analysis on the YFP-CESA6 lines, as described previously (Zhang et al., 2019; 2021b). We found that in wild type, the majority of successful insertion events had an obvious pause phase followed by a steady movement phase, indicating a functional CSC complex; however, in various CESA6 mutation lines, we frequently observed a type of newly-delivered CSC that remained in the stationary phase until the end of the time lapse movie (> 7 min), and this type of behavior was rarely detected for wild-type YFP-CESA6 (Figure 3, A and B). We named this abnormal type of CSC “long-pause” to contrast with the functional particles. Since the mutated amino acids and conserved motifs are hypothesized to have a role in CESA6 catalytic activity, it is likely that the long-pause CSC population represents the delivery and insertion of enzymatically-deficient CSCs that failed to synthesize cellulose microfibrils.

**Figure 3.**
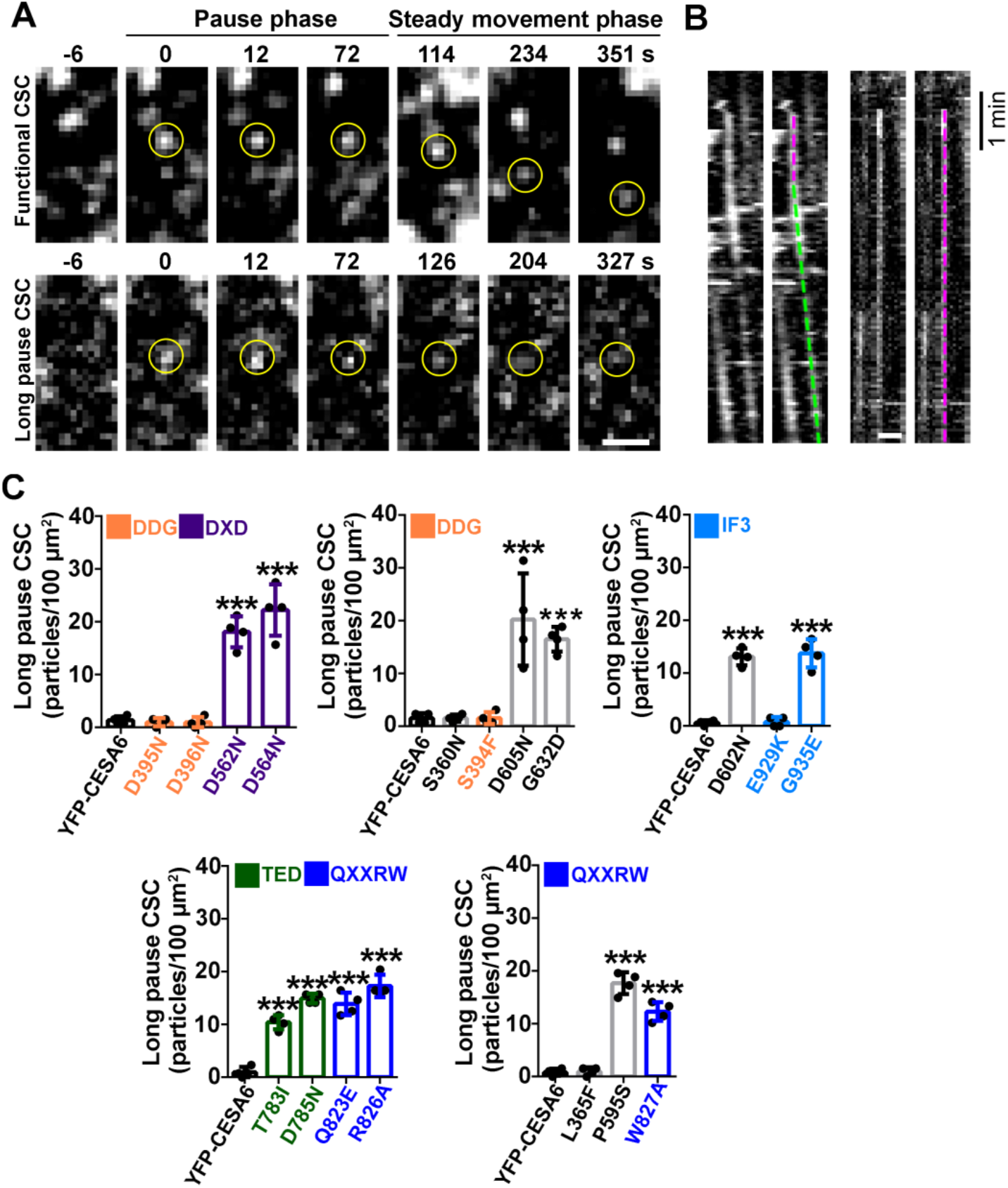
A population of “long-pause” CSCs is caused by mutations in the DXD, TED and QXXRW motifs. **(A)** Representative time-lapse series illustrate single-particle insertion events for a functional CSC and a “long-pause” or nonfunctional CSC. For the functional CSC, after a CSC particle (yellow circle) arrives in the cortex, it pauses (marked as 0 s) for ~ 80 s and exhibits a stationary phase, which is assumed to correspond to tethering, docking, fusion, and delivery of a CSC to the PM, and possibly the activation of CSC. After the functional CSC particle is inserted into the PM, it undergoes steady linear movement as an active complex. For a long-pause CSC, after the particle arrives, it remains stationary at a fixed location for greater than 3 min (85 s + 93 s; mean + 3 SD of pause in wild type) and never shows a translocation phase. Scale bar: 1 μm. **(B)** Representative kymographs of the CSC insertion events for functional and long-pause CSCs as shown in A. For the functional CSC, the duration of the pause phase was determined by fitting a straight line (green) along the moving trajectory and another line (magenta) along the pause phase. The intersection of the two lines was defined as the end of the pause phase. For the long-pause CSC, after arrival at the PM, it fails to translocate and the kymograph only shows a continuous pause phase represented by straight vertical line (magenta). Scale bar: 1 μm. **(C)** Quantification of the density of long-pause CSCs for 18 different mutated CESA6 transgenic lines. Data represent mean ± SD (n = 4 seedlings per genotype, 1 ROI per cell and 2 cells from each seedling were analyzed). *** indicates p < 0.001 by one-way ANOVA followed by Dunnett’s multiple comparisons test compared with wild-type YFP-CESA6 complementation line.

We quantified the density of long-pause CSCs in the AtCESA6 mutation lines, using a method modified from the quantification of CSC tethering frequency, as described previously (Zhang and Staiger, 2022). We extracted all of the kymographs from a region of interest (ROI) at the PM in a 5-min time lapse series and all vertical lines that lasted for at least 3 min were counted as long-pause events. The 3 min cut-off was based on the mean pause time of newly-inserted CSCs in wild type plus 3 SD (85 s + 93 s), so that normal CSC pause events were excluded from the analysis. Our results showed that long-pause CSCs were rarely detected in wild type (< 2 particles/100 μm^2^), whereas a significant increase of long-pause CSCs was detected in some of the mutants, with a range of 10–22 particles per 100 μm^2^ (Figure 3C). Moreover, the increase in long-pause particles correlated with the reduction of mean CSC motility, because all 12 mutations with reduced CSC speed exhibited an increased density of long-pause CSCs, whereas the other six mutations which did not affect CSC speed showed no apparent increase in long-pause CSCs (Figure 2 and Figure 3).

Since five mutants (S360N, L365F, D562N, G632D and E929K) showed reduced *AtCESA6* transcript levels, although AtCESA6 protein abundance in these lines was comparable to wild type (Supplemental Figure S3), we chose a second independent transgenic line for each to confirm the effects of these mutations on CSC motility. The second transgenic lines for these mutants also showed decreased *AtCESA6* transcription levels compared to wild type, which is similar to the original lines (Supplemental Figure S5A). In addition, AtCESA6 protein abundance in these mutants was comparable to wild type (Supplemental Figure S5, B and C). Quantification of CSC speed and long-pause CSC density in these mutants showed that the phenotypes were consistent with those in the original lines (Figure 2, Figure 3 and Supplemental Figure S5).

In total, these results indicate that the DXD, TED, and QXXRW motifs play an important role in enzymatic activity of CESA6 and even a single amino acid mutation in those motifs causes significant defects in CSC motility. In contrast, the DDG motif is likely to be nonessential for CESA catalytic activity, or a single mutation in the DDG motif is not sufficient to cause any measurable effect on motility. The DDG motif is thought to coordinate the donor UDP-glucose molecule at the catalytic site (Purushotham et al., 2020; Daras et al., 2021; Supplemental Table S2).

### The abundance of YFP-CESA6 in the Golgi is increased by mutations in the DDG motif and decreased by mutations in the QXXRW motif

Cellulose synthase proteins are synthesized and translocated into the endoplasmic reticulum (ER) membrane bilayer. After vesicle transport from the ER, CSC rosettes are presumably assembled in the Golgi apparatus and then delivered to the PM via post-Golgi secretory vesicles (McFarlane et al., 2014; Polko and Kieber, 2019; Figure 4A). Previous results show that ES20 treatment results in CSC trafficking defects that include increased CESA6 fluorescent signal in the Golgi as well as an increased abundance of cortical CESA-containing vesicles or SmaCCs, which correlate with reduced CSC delivery rate and reduced CSC density at the PM (Huang et al., 2020). These results suggest that ES20-targeted CESA6 motifs or the catalytic domain play a role in the secretory trafficking of CESA proteins to the PM. We took advantage of the collection of YFP-CESA6 mutant lines and performed high spatiotemporal resolution live-cell imaging to quantitatively assess the dynamics of CESA trafficking through the secretory pathway from Golgi to PM and investigated whether mutations in key motifs alter one or more steps of the endomembrane trafficking process.

**Figure 4.**
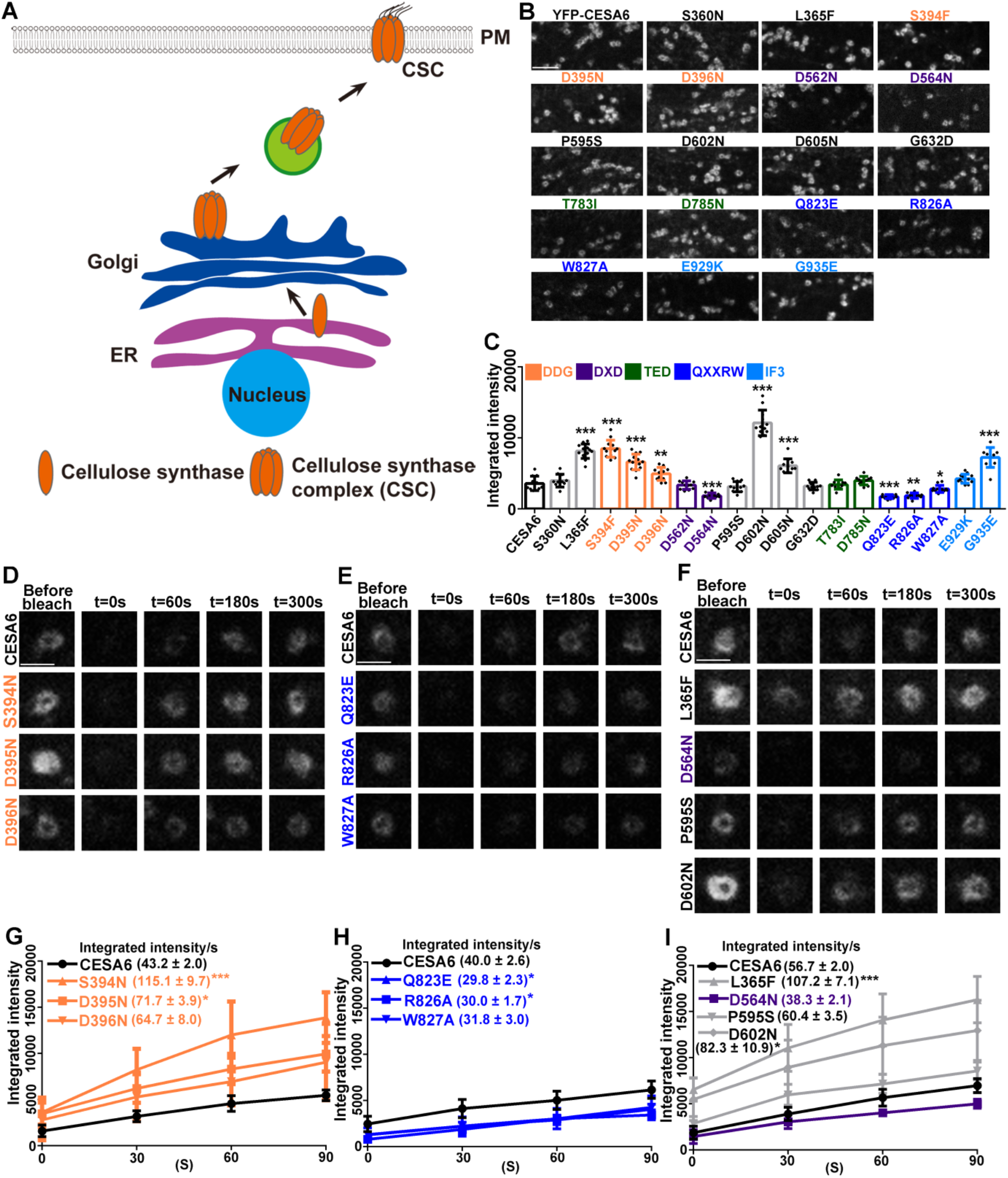
The abundance and recovery of YFP-CESA6 in Golgi are increased and decreased, respectively, by mutations in DDG and QXXRW motifs. **(A)** Cartoon of the cellulose synthase trafficking pathway from ER to PM. Cellulose synthase proteins are synthesized and translocated into the endoplasmic reticulum (ER) membrane bilayer. After vesicle transport from ER, the cellulose synthase complex (CSC) is presumably assembled in the Golgi apparatus and then transported via secretory vesicles to the PM. (**B**) Representative images of YFP-CESA6 at the Golgi in etiolated hypocotyl epidermal cells of transgenic plants expressing wild-type and mutated CESA6. Scale bar: 5 μm. **(C)** Quantification of the fluorescence intensity of YFP-CESA6 at Golgi as described in B. Data represent mean ± SD (n = 12 seedlings, 5 Golgi from each seedling were analyzed). * indicates p < 0.05, ** indicates p < 0.01 and *** indicates p < 0.001 by one-way ANOVA followed by Dunnett’s multiple comparisons test compared with wild-type YFP-CESA6 complementation line. **(D–I)** The recovery of YFP-CESA6 fluorescence in Golgi is increased and decreased, respectively, by mutations in the DDG and QXXRW motifs following photobleaching. **(D–F)** Representative images of Golgi-localized YFP-CESA6 in transgenic plants expressing DDG motif mutations (D), QXXRW motif mutations (E), and four additional CESA6 mutated lines (F) during a fluorescence recovery after photobleaching or FRAP assay. Scale bars: 2 μm. **(G– I)** Recovery of YFP-CESA6 fluorescence at Golgi within 2 min after photobleaching of the seedlings described in D to F, respectively. Data represent mean ± SD (n = 8 seedlings, at last 2 ROIs from each seedling were analyzed). * indicates p < 0.05 and *** indicates p < 0.001 by one-way ANOVA followed by Dunnett’s multiple comparisons test compared with wild-type YFP-CESA6 complementation line.

The localization or abundance of CESAs in the Golgi was investigated first, because Golgi are the earliest location in the forward secretory pathway where YFP-CESA signal can be detected with fluorescence microscopy (Paredez et al., 2006). We detected Golgi-localized YFP-CESA6 signal in each of the 18 transgenic lines; however, there were obvious differences in fluorescence intensity among different mutant lines (Figure 4B). To further evaluate these differences, we quantified the fluorescence intensity of YFP-CESA6 in multiple individual Golgi from several cells and found that seven mutated AtCESA6 lines had significantly increased YFP intensity in the Golgi (Figure 4C), including all three mutations in the DDG motif (S394F, D395N and D396N), the G935E mutation in the IF3 motif, as well as three other mutations not mapped to known motifs (L365F, D602N and D605N). A second independent transgenic line for D602N, with comparable protein and transcript levels to wild type (Supplemental Figure S5), also had increased YFP-CESA6 fluorescence intensity in the Golgi (Supplemental Figure S5, H and I), confirming that this was not due to altered expression of the mutated protein. In contrast, four YFP-CESA6 mutation lines showed decreased Golgi fluorescence intensity, including three mutations in the QXXRW motif (Q823E, R826A and W827A), and D564N in the DXD motif. The seven remaining mutated lines had fluorescence intensities comparable to the wild-type YFP-CESA6 complementation line.

To confirm the specific effects of AtCESA6 mutations on the fluorescence intensities of Golgi-localized YFP-AtCESA6, we generated co-expression lines with two different Golgi markers, mCerulean-MEMB12 and mCherry-SYP32 (Geldner et al., 2009), and tested whether general Golgi protein abundance was generally affected by different CESA6 mutations. As shown in Supplemental Figure S6, the fluorescence intensities of YFP-CESA6 were increased and decreased by DDG and QXXRW motifs mutations, respectively, which is consistent with the original *cesa6* mutants, whereas the fluorescence intensities of mCerulean-MEMB12 and mCherry-SYP32 were not significantly different from wild type, suggesting that mutations of CESA6 do not alter the abundance of other Golgi-localized proteins.

One possibility for the altered CESA6 intensity is that Golgi import/export is altered when certain motifs are mutated. To test this, we utilized a fluorescence recovery after photobleaching (FRAP) assay to analyze the recovery rate of newly-synthesized YFP-CESA6 transported from ER to Golgi. Here, we chose several mutations in the DDG and QXXRW motifs, which showed increased and decreased abundance in the Golgi, respectively, as well as four other mutations for further study. L365F and D602N showed increased abundance in Golgi, whereas D564N and P595S had decreased and comparable abundance of YFP-CESA6 fluorescence in the Golgi, respectively.

Etiolated hypocotyls were treated with 6 μM Latrunculin B (LatB) for 30 min to immobilize the Golgi and individual Golgi were selected for FRAP analysis. As shown in Figure 4, D and G, all three mutations in the DDG motif showed faster recovery of YFP-CESA6 signal compared to wild-type. The S394N and D395N lines, exhibited significantly higher recovery rates and increased by 167% and 66% compared to wild-type YFP-CESA6, respectively. Although D396N had a 50% increase of recovery rate, the difference was not significant. For the QXXRW mutations, all three showed slower recovery of YFP-CESA6 signal compared to wild-type (Figure 4, E and H). The Q823E and R826A lines showed significantly lower recovery rates and were reduced by 26% and 25% compared to wild-type YFP-CESA6, respectively. Although W827A in the QXXRW motif showed a 21% reduction in recovery rate, the difference was not significant. The L365F and D602N mutations, showed significantly higher recovery rates and increased by 89% and 45% compared to wild-type YFP-CESA6, respectively (Figure 4, F and I). The D564N mutation showed 33% reduction in recovery rate, but the difference was not significant, whereas P595S had normal Golgi fluorescence and showed a comparable recovery rate to wild-type YFP-CESA6 (Figure 4, F and I). Taken together, there appears to be a positive correlation between the abundance of YFP-CESA6 in the Golgi and recovery rate in FRAP experiments. These phenotypes are suggestive of defects in CSC rosette formation, abnormal retention in the Golgi cisternae, altered delivery or exit rates, or all of the above.

### Post-Golgi trafficking and secretion of CSCs to the PM is affected by multiple mutations on CESA6

We next investigated whether the post-Golgi trafficking steps were altered by any of the 18 YFP-CESA6 mutations. Since many of the mutations caused major defects in the Golgi, any potential defects in the subsequent Golgi–secretory vesicle–PM trafficking steps could be at least partially or fully due to the altered Golgi phenotype. We applied quantitative live cell imaging tools and measured the density of small CESA-containing compartments (SmaCCs) in the cortical and subcortical cytoplasm, the delivery/insertion rate into the PM, as well as CSC abundance in the PM.

To measure SmaCC abundance, we collected time lapse series of z-stack images and combined 0.2 μm optical sections into cortical (0 to 0.4 μm below the PM) and subcortical (0.6 to 1 μm below the PM) cytoplasm (Sampathkumar et al., 2013; Zhang et al., 2019) and analyzed the density of SmaCCs in both regions for all mutated YFP-CESA6 lines (Figure 5). Similar to our previous analyses (Zhang et al., 2019), the majority of wild-type YFP-CESA6 containing SmaCCs were located in the cortical region, and only L365F and D602N showed increased abundance with ~ 40% and ~ 50% increases, respectively, compared with wild type (Figure 5, A and B). Interestingly, these two mutated CESA6 lines also had significantly higher Golgi abundance and increased FRAP recovery rates (Figure 4, C and I), suggesting that increased abundance of cortical SmaCCs may share the same mechanism as the increased CESA abundance in the Golgi. Two DXD motif mutations (D562N and D564N), P595S, G632D, D785N, and two QXXRW motif mutations (Q823E and R826A) had significantly decreased cortical SmaCC abundance, ranging from 30 to 60% reductions, whereas the remaining nine mutated lines had comparable density to the wild type (Figure 5, A and B). For subcortical SmaCC abundance, only D562N and three QXXRW motif mutations (Q823E, R826A and W827A) had significantly reduced density (Figure 5, C and D). Notably, these mutations also had lower Golgi fluorescence intensity and slower FRAP recovery rates (Figure 4, E and F), suggesting the major defect in trafficking occurs upstream of SmaCC delivery to the PM.

**Figure 5.**
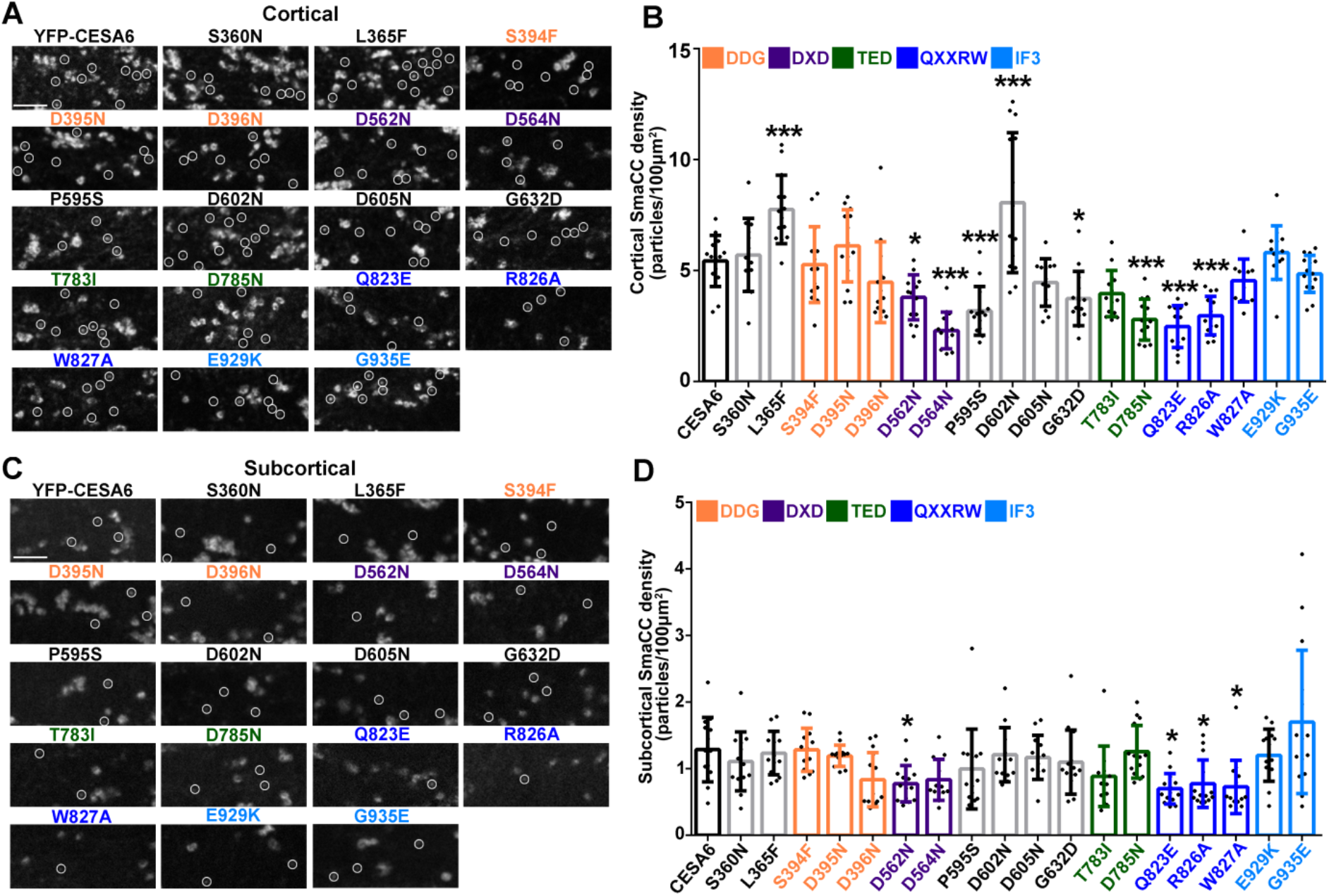
The density of cortical SmaCCs is reduced by mutations in the DXD and QXXRW motifs. **(A, C)** Representative images of SmaCCs in the cortical and subcortical regions of etiolated hypocotyl epidermal cells of transgenic seedlings expressing wild-type and mutated YFP- CESA6. White circles indicate designated SmaCCs. Scale bars: 5 μm. **(B, D)** Cortical and subcortical SmaCC density was measured from images such as those described in A and C. Data represent mean ± SD (n = 12 cells from 12 seedlings, 1 ROI per cell). * indicates p < 0.05 and *** indicates p < 0.001 by one-way ANOVA followed by Dunnett’s multiple comparisons test compared with wild-type YFP-CESA6.

Because at least a subpopulation of the cortical SmaCCs is likely to be involved in secretion of CSCs to the PM, we hypothesized that these YFP-CESA6 mutations would affect the secretion of CSCs to the PM. To analyze bulk secretion rates, we used a FRAP assay to photobleach a region of the cell cortex and counted CSCs that were newly-delivered to this region (Zhang et al., 2019; Huang et al., 2020). Newly-delivered functional CSC particles in the bleached region were defined as those that exhibited a stationary phase followed by steady migration (Li et al., 2016). As shown in Figure 6, A and B, except for the IF3 motif mutation E929K and S360N, the other 16 mutations all showed significantly reduced delivery rates compared with wild-type AtCESA6, and the reductions ranged from ~ 40 to 70%. It should be noted that because only functional CSCs were quantified as new delivery events in the FRAP assay, the delivery rate in mutants that had increased long-pause CSCs may be underestimated.

**Figure 6.**
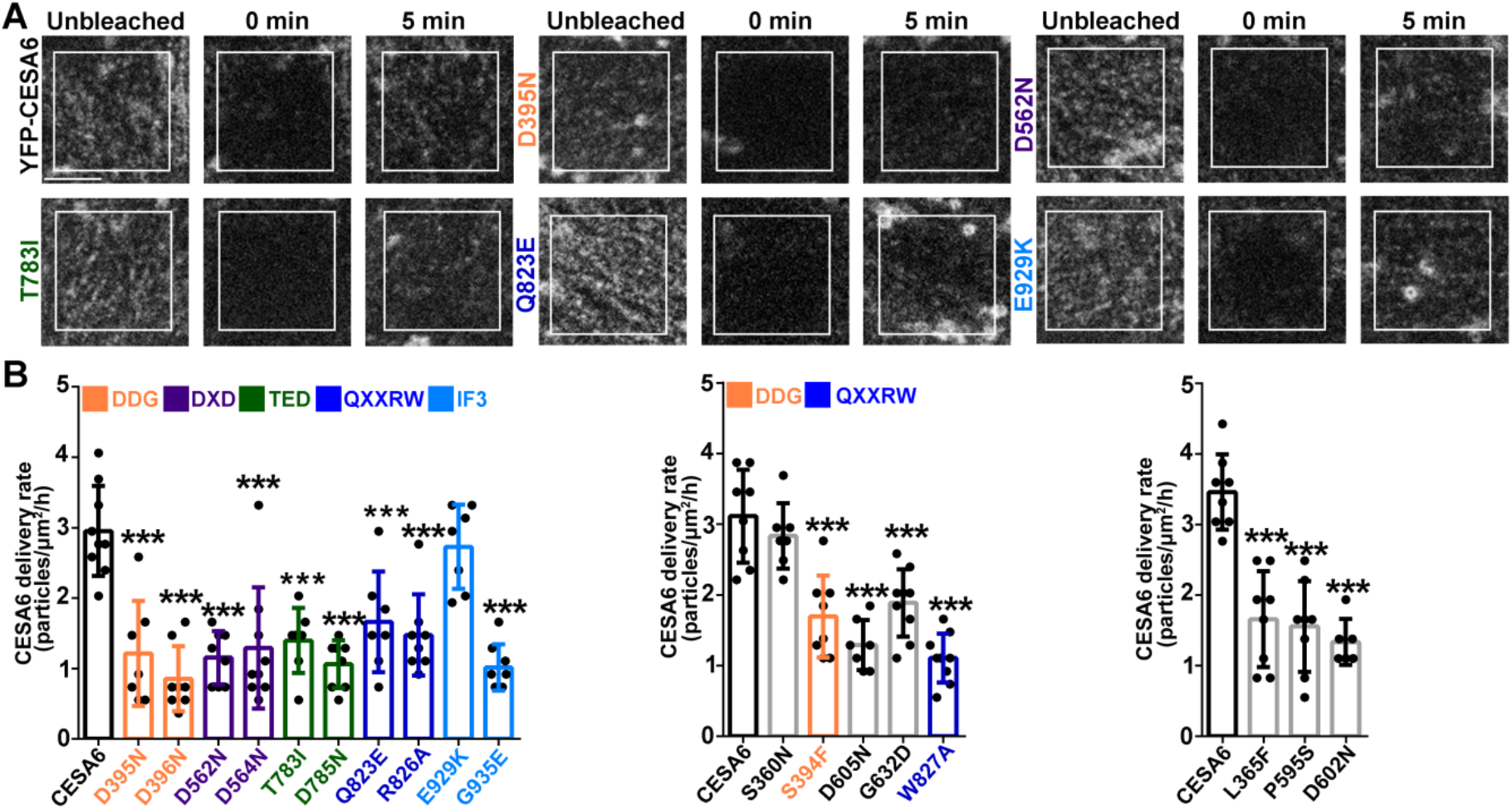
The delivery of functional CSCs into the PM is reduced by mutations in the DDG, DXD, TED and QXXRW motifs. **(A)** Representative images of FRAP analysis for the insertion of functional YFP-CESA6 particles into the PM of etiolated hypocotyl epidermal cells from transgenic seedlings expressing wild-type and mutated YFP-CESA6. Scale bar: 5 μm. **(B)** CSC delivery rates were determined based on the FRAP assays of transgenic seedlings expressing wild-type and mutated YFP-CESA6. Long-pause CSC events were not counted in the CESA6 delivery rate quantification. Data represent mean ± SD (n ≥ 7 seedlings, 2 ROIs from each seedling were analyzed). *** indicates p < 0.001 by one-way ANOVA followed by Dunnett’s multiple comparisons test compared with wild-type YFP-CESA6 complementation line.

To understand whether these mutations also affect the endocytosis of CSCs, we treated the 18 mutants with Brefeldin A (BFA), a fungal toxin, which inhibits vesicle trafficking and caused the accumulation of endosome and TGN into endomembrane compartments named “BFA bodies” (Nebenfuhr et al., 2002). BFA treatment blocks the recycling of endocytosis-related SmaCCs to the PM and hence causes the accumulation of SmaCCs (Gutierrez et al., 2009). After 50 μM BFA treatment for 2 h, we found visible accumulation of cortical SmaCCs in wild type and all of the mutants except S360N (Supplemental Figure S7, A to D); however, the increased ratio (BFA/DMSO) of cortical SmaCCs in these mutants was comparable to wild type (Supplemental Figure S7, E to G). For subcortical SmaCCs, BFA treatment did not cause obvious accumulation in either wild type or any mutants (Supplemental Figure S7, H to K). These experiments indicate that mutations in key motifs of the CESA6 catalytic domain did not alter the endocytosis of CSCs.

The abundance of CSCs at the PM could influence the content of cellulose in the cell wall; therefore, we measured the density of CSCs in the PM for wild type and all mutated YFP- CESA6 lines. As shown in Figure 7, A and B, only D564N and Q823E exhibited significantly increased CSC density of 35% and 25%, respectively, compared with wild type. In contrast, six mutations including L365F, DDG motif mutations (S394F and D395N), D602N, and IF3 motif mutations (E929K and G935E) showed significantly decreased PM density and the remaining ten lines had CSC density comparable to wild type (Figure 7, A and B).

**Figure 7.**
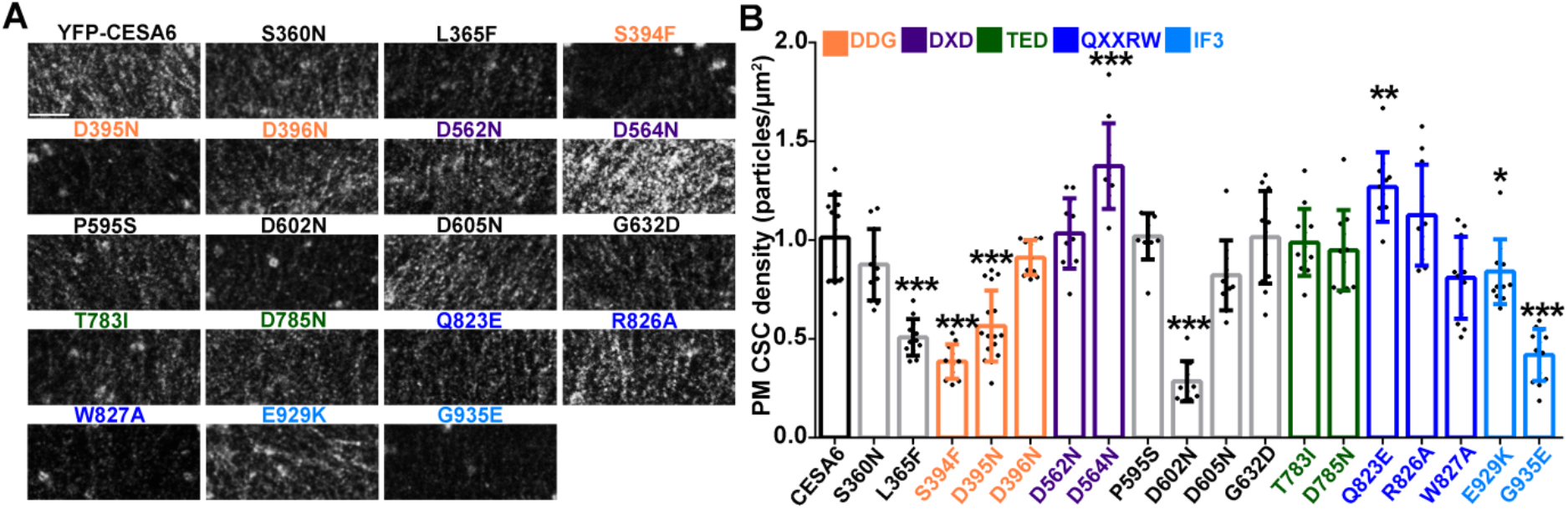
The density of CSCs at the PM is differentially affected by numerous mutations in the CESA6 catalytic domain. **(A)** Representative single timepoint images of CSCs at the PM in etiolated hypocotyl epidermal cells expressing wild-type and mutated YFP-CESA6. Scale bar: 5 μm. **(B)** CSC density at the PM. Data represent mean ± SD (n = 12 seedlings, 1 ROI per cell and 2 cells from each seedling were analyzed). * indicates p < 0.05, ** indicates p < 0.01 and *** indicates p < 0.001 by oneway ANOVA followed by Dunnett’s multiple comparisons test compared with wild-type YFP- CESA6.

### The abundance of YFP-CESA6 in the Golgi positively correlates with cortical SmaCC density and negatively correlates with cellulose content and abundance of CSCs in the PM

Overall, we quantified crystalline cellulose content, seedling growth, CSC motility and subcellular trafficking dynamics in 18 mutated YFP-CESA6 complementation lines and the results are summarized in Supplemental Table S3. With this large data set, we performed pairwise correlation analyses to determine whether there was any relationship between cellulose biosynthesis and other parameters in the mutant lines. For correlation analyses, the actual values for each parameter were normalized to wild-type YFP-CESA6 and expressed as a percentage (Supplemental Table S3). As shown in Supplemental Figure S8, cellulose content positively correlated with hypocotyl length and negatively correlated with cortical SmaCC density and CSC Golgi intensity. Surprisingly, there was no obvious correlation between crystalline cellulose content and the rate of CSC motility, although we found that mutations in most key motifs of the catalytic domain of CESA6 showed reduced CSC speed. Similarly, long-pause CSC density, CSC Golgi recovery rate, Golgi ring diameter, CSC delivery rate and CSC PM density also did not show obvious correlations with cellulose levels.

Since the abundance of YFP-CESA6 in the Golgi was the earliest phenotype detected in the secretory pathway (Figure 4), we examined whether there was any correlation between the Golgi and post-Golgi phenotypes measured in the mutant lines by pairwise correlation analysis. As shown in Figure 8, YFP-CESA6 fluorescence intensity in the Golgi positively correlated with fluorescence recovery rate as well as the density of cortical SmaCCs, whereas it negatively correlated with CSC PM density and crystalline cellulose content. There was no obvious correlation between CSC Golgi intensity and other parameters including hypocotyl length, CSC motility, long-pause CSCs, Golgi ring diameter or CSC delivery rate.

**Figure 8.**
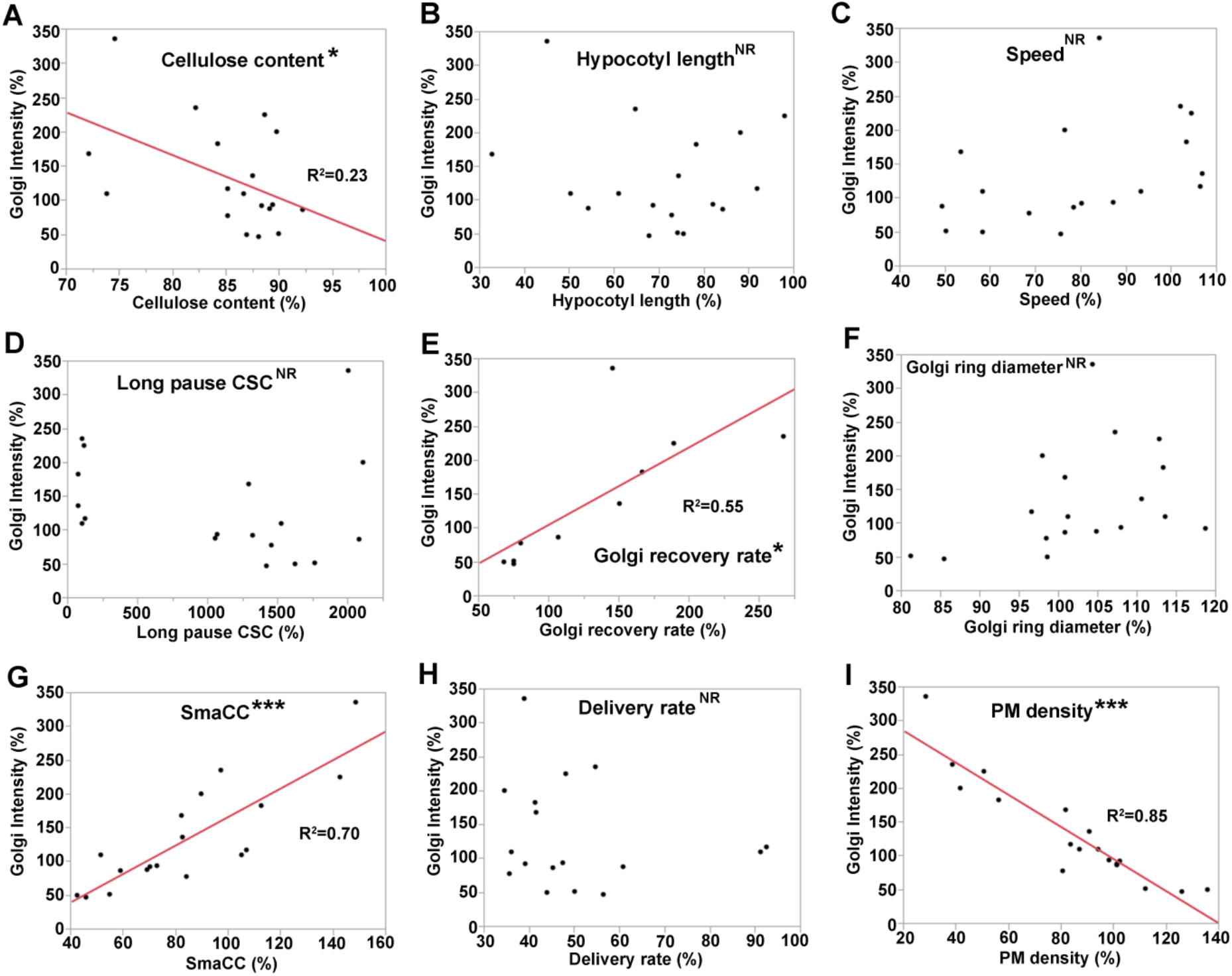
The fluorescence intensity of YFP-CESA6 in the Golgi positively correlates with Golgi fluorescence recovery rate as well as cortical SmaCC density, and negatively correlates with cellulose content and CSC PM density. **(A-I)** Pairwise correlation analysis between the fluorescence intensity of YFP-CESA6 in the Golgi and cellulose content, hypocotyl length or various CSC trafficking parameters was conducted using JMP 11 software. Fluorescence intensity of YFP-CESA6 in the Golgi was plotted as a function of cellulose content **(A)**, hypocotyl length **(B)**, CSC speed **(C)**, long-pause CSC abundance **(D)**, fluorescence recovery rate of YFP-CESA6 in Golgi **(E)**, Golgi ring diameter (**F**), cortical SmaCC density **(G)**, CSC delivery rate **(H)** and CSC PM density **(I)**. Corresponding parameters in **(A–H)** were collected from 18 YFP-CESA6 mutated complementation lines and Golgi fluorescence recovery rate in **(E)** was obtained from 10 YFP- CESA6 mutated complementation lines. Each parameter was normalized to wild-type YFP- CESA6 and the corresponding relative values were used for the correlation analysis. NR, no predictive relationship; *, P ≤ 0.05, ***, P ≤ 0.001, Bivariate fit/ANOVA for all data points for each parameter.

To explore whether any CSC trafficking parameter explains the most variance of the data, we performed principal component analysis (PCA) on the whole dataset. We found principal component 1 (PC1) and principal component 2 (PC2) accounted for the vast majority of the variation in the data and represented 42% and 23% of variance, respectively (Supplemental Figure S9). PC1 scaled positively with cortical SmaCC density and YFP-CESA6 Golgi fluorescence, but negatively with the density of CSCs at the PM (Supplemental Figure S9). This result suggested that CESA6 mutants with positive PC1 score exhibited high cortical SmaCC density, high YFP-CESA6 abundance in the Golgi and low CSC PM density, and vice versa. PC2 scaled positively with hypocotyl length, crystalline cellulose content and CSC speed (Supplemental Figure S9), suggesting mutants with positive PC2 score exhibited high cellulose content, increased hypocotyl length and high CSC speed, and vice versa.

Based on the pairwise correlation (Figure 8 and Supplemental Figure S8) and PCA analyses (Supplemental Figure S9), as well as the distinct and well-correlated Golgi–SmaCC–PM trafficking phenotypes, we found that the 18 YFP-CESA6 mutant lines can be separated into two major groups (Supplemental Table S3, Supplemental Figure S9C). Group I comprises nine mutations including S360N, S394F, D395N and D396N in or near the DDG motif, E929K and G935E in the IF3 motif, and three other mutations L365F, D602N and D605N. Except for S360N and E929K, the remaining seven mutations caused significantly increased Golgi intensity. In addition, most of the Group I mutations showed significantly reduced CSC density at the PM, with either an increase or no significant change in the density of cortical SmaCCs compared with wild-type. Group II mutations had the opposite phenotypes: significantly decreased Golgi intensity and/or SmaCC density, whereas the PM CSC density was either increased or not significantly changed compared with wild-type. Nine mutations fall into this group: D562N, D564N, P595S, G632D, T783I, D785N, Q823E, R826A and W827A. Strikingly, all mutations in Group II caused significantly reduced CSC speed at the PM, with seven of the nine belonging to the DXD, TED and QXXRW motifs, whereas in Group I, six of the nine mutations had no significant effect on CSC motility, including the DDG motif. The distinct patterns of Golgi-to-PM phenotypes between Group I and II are perhaps due to two different types of mechanisms; the former is likely associated with CSC trafficking defects, whereas the latter is more likely due to protein misfolding or rosette formation defects in the ER or Golgi. When the homologous single amino acid mutations were mapped on the recently solved AtCESA3 catalytic domain structure (Qiao et al., 2021), most Group II mutations motifs were close to the homodimer interface formed by β-strand 6 and α-helix 9, whereas Group I mutations were distal to the homodimer interface (Supplemental Figure S10).

### CSC assembly in the Golgi is affected by mutations in QXXRW motif

To investigate whether Group I or II mutations altered CESA complex formation in Golgi, we first analyzed the diameter of YFP-CESA6 ring structure in Golgi. A previous study of the Golgi localized CESA-interacting STELLO proteins shows that a *stl1stl2* double mutant has decreased YFP-CESA6 ring diameter as well as a CSC assembly defect (Zhang et al., 2016). Our results showed that two Group II mutations, Q823E and R826A from the QXXRW motif, had a decreased Golgi ring diameter (Figure 9, A and B). The ring diameter reduction of YFP- CESA6 in Golgi may be related to a CSC assembly defect. To further test this hypothesis, we extracted microsomes from various YFP-CESA6 mutation lines and performed blue-native polyacrylamide gel electrophoresis (BN-PAGE). The CESA complexes could be detected as a multimeric protein complex by BN-PAGE (Figure 9, C and H), as a single band at ~ 480 kDa which is likely to be a trimeric complex and is consistent with previous studies (Wang et al., 2008; Zhang et al., 2016; Wen et al., 2022). YFP-CESA6 band intensity in the complex was significantly decreased in Q823E and R826A lines, but not in other tested mutants, when compared with wild type (Figure 9, D and I). When YFP-CESA6 band intensity in the complex was normalized against total YFP-CESA6 protein abundance analyzed by SDS-PAGE (Figure 9, C to L) the results confirmed that there was significantly reduced CSC complex formation in Q823E and R826A mutation lines. The decreased YFP-CESA6 ring diameter and reduced abundance of CESA complex in Q823E and R826A mutations indicate that the QXXRW motif is likely involved in CSC assembly.

**Figure 9.**
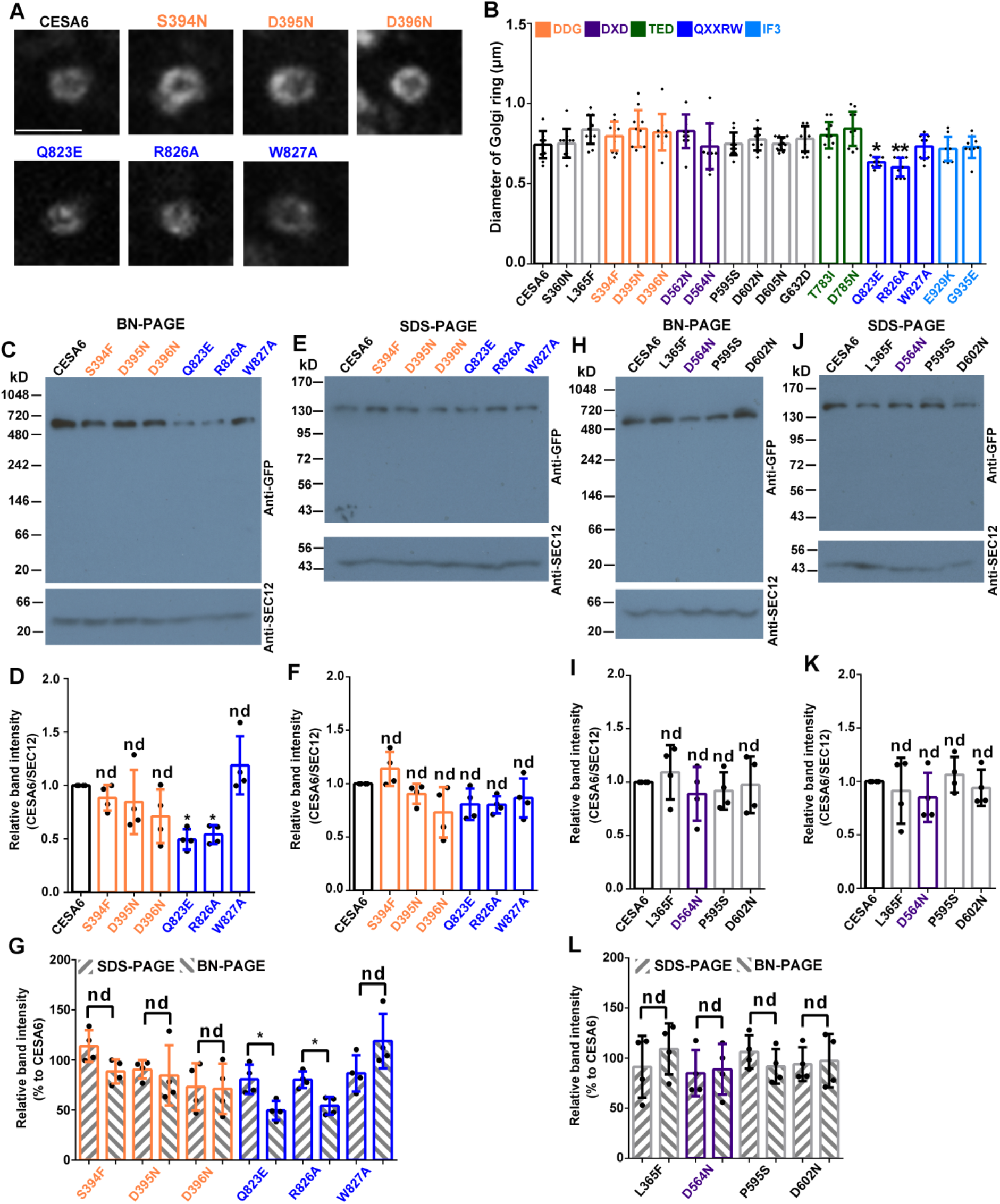
CSC complex formation is reduced by mutations in the QXXRW motif. **(A)** Representative images of Golgi in etiolated hypocotyl epidermal cells of transgenic plants expressing wild-type and mutated YFP-CESA6. Scale bar: 2 μm. **(B)** Measurement of the fluorescent Golgi ring diameter from transgenic seedlings expressing wild-type and mutated YFP-CESA6. Data represent mean ± SD (n = 10 seedlings, 5 Golgi from each seedling were chosen for analysis). * indicates p < 0.05 and ** indicates p < 0.01 by one-way ANOVA followed by Dunnett’s multiple comparisons test compared with wild-type YFP-CESA6 complementation line. (**C–L**) CSC complex formation is affected by two of three mutations in the QXXRW motif. (**C** and **H**) Representative protein immunoblots of Blue Native (BN)-PAGE gels showing the abundance of YFP-CESA6 and SEC12 in microsomes isolated from seedlings of wild type and various CESA6 mutated lines. (**E** and **J**) Representative protein immunoblots from denaturing SDS-PAGE gels showing the abundance of YFP-CESA6 and SEC12 in total protein extracts from seedlings of wild type and various CESA6 mutated lines. (**D** and **I**) Quantification of the relative abundance of YFP-CESA6 in microsomes extracted from transgenic lines as shown in C and H. (**F** and **K**) Quantification of the relative abundance of YFP-CESA6 in total protein extracted from the transgenic lines as shown in E and J. (**G** and **L**) Relative band intensity comparison of YFP-CESA6 (normalized with wild type) in BN- PAGE and SDS-PAGE. Data represent mean ± SD (n = 4 biological replicates). Significant differences in D to K were analyzed by one-way ANOVA followed by Dunnett’s multiple comparisons test compared with wild-type YFP-CESA6. * indicates p < 0.05, nd indicates no significant difference. Significant differences in G and L were analyzed by Student’s *t*-test. * indicates p < 0.05, nd indicates no significant difference.

## Discussion

Understanding the molecular basis of cellulose biosynthesis is a key question for plant biology, not only because of its biological importance, but also due to the great economic potential of cellulose. Cellulose biosynthesis inhibitors (CBIs) combined with chemical genetics are powerful tools to dissect the molecular mechanisms of cellulose biosynthesis and CESA protein function (Tateno et al., 2016; Huang et al., 2020; Larson and McFarlane, 2021). The recently resolved cryo-EM and crystal structures of angiosperm CESAs also greatly enhance our understanding of the molecular basis of cellulose biosynthesis and CSC rosette formation (Purushotham et al., 2020; Qiao et al., 2021; Zhang et al., 2021c). In this work, guided by the previously characterized *es20r* point mutation collection as well as the predicted ES20-binding site on CESA6 (Huang et al., 2020), we provide a comprehensive mutational analysis of the conserved CESA6 catalytic domain and explored its roles in catalytic activity, CESA endomembrane trafficking and protein dynamics regulation.

All 18 AtCESA6 mutations created here affected cellulose biosynthesis (Figure 1E, Supplemental Table S3), and were predicted to interfere with catalytic activity by disrupting UDP-glucose binding, glucan chain elongation, or glucan chain translocation (Supplemental Table S2). Briefly, DDG as well as DXD mutations together with the surrounding mutations including S360N and L365F are predicted to affect UDP-glucose coordination; the TED and QXXRW motif mutations could affect glucose addition to the elongating glucan chain and hence interfere with glucan chain elongation; IF3 mutations E929K and G935E may interfere with glucan chain translocation. However, the trafficking defects observed in the YFP- AtCESA6 single amino acid replacement mutants could be explained by altered Golgi import/export, improper protein folding, defective CSC rosette formation, or any combination of these, in addition to effects on catalytic activity.

The catalytic activity of CESA elongates and extrudes glucan chains into the cell wall, and this polymerization is argued to be the driving force for CSC movement in the plane of the PM (Paredez et al., 2006). Here, we used a battery of mutations in and around the catalytic core of CESA6 to test this hypothesis. CBIs like ES20 and ES20-1 that bind at the catalytic site and presumably reduce catalytic activity also significantly reduce CSC motility and offer indirect support for this model (Huang et al., 2020; 2021). In this study, single mutations in DXD, TED and QXXRW motifs caused significantly reduced CSC speed as well as a marked increase in the abundance of long-pause CSCs at the PM. These results support the hypothesis that PM- localized CSC motility correlates with catalytic activity. By contrast, three mutations in the DDG motif showed comparable speed to wild-type and no increase in long-pause CSCs, although they all showed a reduction in crystalline cellulose content. In addition, S360N and L365F, close to the DDG motif in the 3D structure of CESA protein, did not affect CSC motility or the abundance of long-pause CSCs. These results suggest that the DDG motif, together with the adjacent S360 and L365, may not be essential for catalytic activity. A previous report showed that mutations in the DDG motif of secondary cell wall related CESAs complemented cellulose biosynthesis and plant growth better than mutations in DXD and TED motifs when introduced into the null mutants (Kumar et al., 2018). Based on the 3D structures of RsBcsA and PttCESA8, although DDG and DXD motifs are both involved in the coordination of UDP-glucose, DDG is distal to the core catalytic region whereas the DXD motif is proximal (Morgan et al., 2013; Purushotham et al., 2020), implying that the distance from the core catalytic region plays a significant role..

In addition to these key motifs, several other mutations including P595S, D602N, D605N and G632D also caused significant reductions in CSC motility and increased the abundance of long-pause CSCs. The conserved P595 is located in a previously undescribed motif near the catalytic core (Daras et al., 2021) and mutations of this region in other CESA isoforms show similar phenotypes compared to AtCESA6^P595S^, originally described as *es20r10* (Huang et al., 2020). For example, analogous mutations of AtCESA3 (*repp3/thanatos*, AtCESA3^P578L/S^) and AtCESA7 (*fra5*, AtCESA7^P557S^) cause severe growth defects and cellulose biosynthesis deficiencies (Zhong et al., 2003; Daras et al., 2009; Feraru et al., 2011). In addition, a previous report showed *any1* (AtCESA1^D604N^), the analogous mutation to AtCESA6^D605N^, caused a reduction in CSC speed (Fujita et al., 2013). Similarly, *rsw1-2* (AtCESA1^G631S^), the analogous mutation to AtCESA6^G632D^, caused radial root swelling and cellulose content reduction (Gillmor et al., 2002). Interestingly, P595, D602, D605 and G632 are flanked by the conserved PCR and CSR domains and three of them (P595, D605 and G632) are conserved between plant and bacterial CESAs (Huang et al., 2020). In addition, these four amino acids are close to DXD, TED and QXXRW motifs in the 3D structures (Figure 1, A and B), indicating they could participate in UDP-glucose binding, glucan chain initiation and elongation. Nonetheless, P595S and G632D are classified as Group II mutations mimicking disruptions to the QXXRW motif, whereas D602 and D605 are Group I mutations resembling those in the DDG motif (Supplemental Table S3). These results suggest that the region flanked by PCR and CSR domains may be implicated in catalytic activity, protein folding, CSC rosette formation, or vesicle trafficking.

The interfacial helix3 (IF3) packs near the catalytic domain, according to the crystallographic structure of RsBcsA and the cryo-EM structure of PttCESA8 and may be involved in glucan chain translocation (Morgan et al., 2013; Purushotham et al., 2020). Consistent with this, the G935E mutation in IF3 caused the occurrence of long-pause CSCs and slowed CSC speed whereas another IF3 mutation, E929K, did not, although both mutations affected crystalline cellulose levels. Our results suggest that G935E likely interfered with glucan chain translocation and thereby affected CSC movement in the PM. Surprisingly, E929 faces the glucan translocation channel formed by transmembrane domains in the predicted structure of CESA6 (Figure 1B), whereas G935 is distal to the channel. Further investigation is needed to determine how these mutations affect glucan chain translocation or CSC catalytic activity.

A major focus of this work was to investigate potential links between the CESA catalytic domain and regulation of CSC trafficking. Because the Golgi apparatus represents the earliest step in the secretory pathway where YFP-CESA is visible, and both pairwise correlation and PCA analysis revealed that the Golgi fluorescence intensity phenotype in YFP-CESA6 mutants accounted for a majority of the variation in the data, we propose that Golgi is the primary location where mutation-induced defects occur and the downstream post-Golgi trafficking phenotypes are a consequence of the Golgi defects. In addition, the 18 mutants can be categorized into two major groups (Supplemental Table S3) and their differential Golgi phenotypes suggest distinct mechanisms. The Group I defects, including those associated with DDG mutations as well as S360N, L365F, D602N, D605N, E929K, and G935E, are most likely caused by inhibition of Golgi export and/or post-Golgi delivery of CSCs to the PM, resulting in abnormal retention of CESA6 in the Golgi and/or SmaCCs, and thereby reducing the CSC delivery rate and CSC abundance in the PM. Abnormal accumulation of cortical CESA compartments combined with reduced PM delivery has been related to defects in CSC trafficking/secretion (Zhang et al., 2019; Zhang et al., 2021b). Interestingly, Group I mutations are mostly located distal to the CESA catalytic core on the predicted 3D structure of AtCESA6 (Huang et al., 2020; Figure 1B). It is possible that these amino acids are involved in CSC trafficking regulation and/or association with trafficking-related proteins.

In contrast to Group I, the reduction of YFP-CESA6 fluorescence intensity in the Golgi and/or SmaCCs in Group II mutants is likely caused by protein misfolding and/or defective CSC formation in the ER or Golgi. Our results with two mutations in the QXXRW motif that reduced CSC complex formation in the Golgi support this hypothesis. All Group II motifs are predicted to reduce CESA catalytic activity and are located in or proximal to the predicted CESA6 catalytic core (Huang et al., 2020; Figure 1B), suggesting that the catalytic core may contain important structural information that is either related to correct protein folding or CSC rosette assembly. The cryo-EM structure of PttCESA8 and negative-staining EM structure of AtCESA1 catalytic domain reveal that the PCR domain participates in CESA trimerization (Purushotham et al., 2020; Du et al., 2022), but none of our 18 mutations were located in the PCR domain, suggesting that these mutations may not directly participate in CSC rosette assembly. So far there is little information on how conserved catalytic core motifs affect CESA structure or rosette formation. Notably, loss of the Golgi-localized, CESA-interacting STELLO proteins disrupts CSC assembly in the Golgi by an unknown mechanism and causes CSC post-Golgi secretion defects (Zhang et al., 2016), suggesting that defects in the Golgi may affect post-Golgi CSC trafficking.

A recent crystal structure of the catalytic domain of the primary cell wall AtCESA3 suggests that the catalytic core is capable of forming a homodimer at an interface involving β-strand 6 (β6) and α-helix 9 (α9) and that homodimerization may play a role in preventing premature CSC assembly (Qiao et al., 2021). Although none of our mutations were analogous to the two predicted residues required for homodimerization in CESA3 (Q571 and C573), most Group II amino acids especially QXXRW motifs are proximal to the homodimer interface formed by β6 and α9, whereas Group I amino acids including S360, L365 and the DDG motif are distal to the homodimer interface (Supplemental Figure S10). It is possible that certain Group II mutations, like Q823E and R826 have an allosteric effect on β6 and α9 and thereby hinder CESA homodimerization and subsequent CSC assembly. Moreover, although all Group II mutations resulted in significantly reduced CSC motility, the extent of Golgi intensity changes did not show significant correlation with CSC speed at the PM (Figure 8), indicating that the Golgi and post-Golgi delivery defects were unlikely related to catalytic activity defects, nor caused by feedback from reduced cellulose synthesis at the cell surface.

Finally, Group I but not Group II mutations phenocopied the ES20-induced trafficking defects including significantly increased YFP-CESA6 abundance in the Golgi, increased cortical SmaCC density as well as reduced rate of CSC delivery and density at the PM (Huang et al., 2020). Perhaps acute treatments with ES20 cause immediate trafficking inhibition of CSCs thereby mimicking the Group I phenotypes but do not induce the potential protein folding or CSC rosette defects similar to Group II mutations. Similarly, the well-studied CBI isoxaben targets CESAs directly and causes rapid trafficking inhibition of CSCs to the PM (Tateno et al., 2016).

Another interesting but controversial correlation highlighted by our data is the positive relationship between crystalline cellulose content in the cell wall and cell expansion or growth (Supplemental Figure S8). Cell expansion is driven by internal forces created through turgor pressure (Cosgrove, 2005; 2016), whereas cell wall biomass organization, primarily through the cellulose microfibrils, resists the forces of turgor pressure. An interesting study shows that cell expansion and cellulose synthesis are independent processes and regulated by different mechanisms (Ivakov et al., 2017). Cell expansion is regulated by protein-mediated changes in cell wall extensibility which is driven by the circadian clock (Yazdanbakhsh et al., 2011; Apelt et al., 2015; Ivakov et al., 2017). By contrast, cellulose biosynthesis is controlled by intracellular CSC trafficking and is independent of the circadian clock (Ivakov et al., 2017). Plant growth requires cellulose for the deposition of new cell wall materials; on the other hand, the network of aligned cellulose microfibrils restricts cell expansion and hence restricts plant growth (Cosgrove, 2005; 2016). Several cellulose biosynthesis mutants show growth defects; for example, *prc1-1/cesa6* (Fagard et al., 2000), *rsw1/cesa1* (Arioli et al., 1998), and *any1/cesa1* (Fujita et al., 2013) all show anisotropic cell swelling. But this is not always the case, for example, the growth phenotype of *prc1-1* could be partially rescued by simultaneous mutation of THESEUS1 (THE1), a receptor-like kinase (Hematy et al., 2007). The *prc1-1 the1* double mutant has comparable levels of cellulose compared with the *prc1-1* single mutant (Hematy et al., 2007), suggesting reduced cellulose content is not the major factor impeding the growth of *prc1-1*. In our study, L365F, P595S, E929K and G935E mutations showed a cellulose biosynthesis deficiency but exhibited normal hypocotyl growth compared to wildtype. Here, we found that most CESA6 mutated lines showed hypocotyl growth defects including mutations in DDG, DXD, TED and QXXRW motifs, and these mutants also exhibited cellulose biosynthesis deficiency.

In a previous publication, D395N, a mutation in the DDG motif of AtCESA6, completely failed to rescue the *prc1-1* hypocotyl growth phenotype, YFP-CESA6 disappeared from the PM and Golgi, and localized instead to ER-like structures (Park et al., 2019). Another mutation, Q823E in the QXXRW motif, completely rescued the *prc1-1* hypocotyl growth phenotype and showed comparable cellulose content and CSC motility compared with wild type (Park et al., 2019). Both findings are inconsistent with our results. Here, we found that D395N and Q823E only partially rescued the *prc1-1* hypocotyl growth phenotype and both showed a reduction in cellulose content. In addition, YFP-CESA6^D395N^ localized to the Golgi, SmaCCs and PM and was categorized as a Group I mutant. The discrepancy between our data and the previous report could be attributed to different CESA6 complementation strategies. We used the CESA6 genomic sequence for complementation, whereas the Park *et al*., 2019 study used the CESA6 coding sequence.

In summary, we comprehensively analyzed the function of the CESA6 catalytic core domain in plant growth, cellulose biosynthesis, and trafficking to the PM. Our results reveal that the CESA catalytic domain, in addition to its enzymatic function, plays a major role in trafficking regulation and/or CSC rosette formation in the ER or Golgi. These data advance our understanding of the cellular and molecular mechanisms underlying cellulose biosynthesis and inform potential engineering strategies to manipulate cellulose production.

## Materials and methods

### Vector construction for YFP-CESA6 single amino acid replacement mutations and generation of transgenic Arabidopsis plants

The wild-type *YFP-CESA6* binary vector containing *CESA6* native promoter (2245 bp), *YFP* coding sequence and *CESA6* genomic sequence with native terminator was created using Gibson assembly assay, as described previously (Huang et al., 2020). The *YFP-CESA6* construct was used as a template to prepare 18 *YFP-CESA6* mutated constructs using a Q5 Site Directed Mutagenesis Kit (New England Biolabs, Ipswich, MA) with the specific primers listed in Supplemental Table S4. All constructs were verified by DNA sequencing and introduced into an Arabidopsis (*Arabidopsis thaliana) prc1-1/cesa6* homozygous mutant line using *Agrobacterium tumefaciens-mediated* floral dip transformation (Clough and Bent, 1998).

### Plant materials, growth conditions, and growth assays

All plants were grown in a Percival growth chamber at 22°C, under light or dark conditions as specified in each experiment. To quantify the hypocotyl length of YFP-CESA6 single amino acid replacement mutants, seeds were sequentially sterilized with 50% bleach and 75% ethanol. After washing with sterile water, seeds were sown on ½-strength Murashige and Skoog (MS) medium with 1% sucrose (for light-grown seedlings) or without sucrose (for dark-grown seedlings) and stratified for 2 d at 4°C. Plates were wrapped in two layers of aluminum foil and placed vertically in the growth chamber at 22°C for 7 d. Seedlings on plates were scanned using an Epson Perfection V550 scanner and hypocotyl length was measured using ImageJ. All experiments were conducted with at least 3 biological repeats.

### Crystalline cellulose content measurements

Seeds of YFP-CESA6 mutants were sown on ½-strength MS medium without sucrose. After stratification for 2 d, plants were grown in the dark for 7 d. The whole hypocotyl tissue without seed coat was used for cellulose measurement. Cell wall preparations and crystalline cellulose measurements were performed as described previously (Huang et al., 2020).

### RNA extraction and RT-qPCR

Total RNA was extracted from 6-day-old light-grown seedlings using TRIzol reagent (Invitrogen, USA) according to the manufacturer’s protocol. RNA (0.5 μg) was reverse transcribed by the PrimeScript RT reagent kit with genomic DNA Eraser (Takara, USA) using random and oligo-dT primer mix in a 10 μL reaction. Bio-Rad SsoAdvanced SYBR Green Supermix (Bio-Rad, USA) was used to perform the RT-qPCR on a Roche LightCycler 96 thermocycler (Roche, USA). The relative gene expression levels were calculated using the 2^-ΔΔCT^ method as described (Livak and Schmittgen, 2001). *AtActin1* was used as an internal control for gene expression normalization. Primers used for RT-qPCR are listed in Supplemental Table S4.

### Protein immunoblot analysis of YFP-CESA6 abundance in transgenic lines

To quantify CESA6 protein levels in different transgenic lines expressing mutated YFP-CESA6 in *prc1-1*, total protein was isolated from 6-d-old light-grown seedlings. The seedlings were ground into a powder under liquid nitrogen and homogenized with lysis buffer (50 mM Tris-HCl, pH 7.5, 150 mM NaCl, 0.5% Triton X-100, 2 mM DTT) with EDTA-free protease inhibitor (Thermo Fisher, USA) at 1:1 ratio (1 mL buffer: 1 g tissue). Homogenized samples were transferred to a 1.5 mL microcentrifuge tube and centrifuged for 30 min at 20,000 *g*, 4°C. The supernatant was collected after centrifugation and saved as the total protein extract.

Protein concentrations of the different transgenic lines were measured by Bradford assay and adjusted to the same levels. Total protein samples were loaded onto SDS-PAGE gels and transferred to nitrocellulose membrane for immunoblot analysis. Anti-GFP (monoclonal antibody raised in mouse, Takara, USA, catalog # 632381) and anti-SEC12 (Bar-Peled and Raikhel, 1997) antibodies were used to detect CESA6 and SEC12. HRP-conjugated secondary antibodies diluted by 1:10000 (HRP-conjugated Goat-anti-Mouse IgG secondary antibody for GFP, Invitrogen, USA, product #31444; HRP-conjugated Goat-anti-Rabbit IgG secondary antibody for SEC12, Invitrogen, USA, product #31460) and SuperSignal West Pico Plus Chemiluminescent Substrate kit (Thermo Scientific, USA) reagent were used for protein band detection. The immunoblot results were detected using X-ray film. To measure the intensity of each protein band, a rectangular box was drawn around each band and the integrated intensity in each box was measured with ImageJ. A box with identical shape and size was used to measure the integrated intensity of background. The background intensity was subtracted from each protein band to obtain the real intensity for each protein band. The CESA6 abundance was normalized against SEC12 abundance for each protein sample by calculating the ratio of CESA6 intensity/SEC12 intensity. Then, the value for each lane was normalized against wildtype YFP-CESA6.

### Microsome extraction and BN-PAGE

Extraction of a microsome fraction and BN-PAGE were performed according to published methods (Wang et al., 2008; Zhang et al., 2016; Wen et al., 2022). Briefly, 14-d-old light-grown seedlings were used for protein extraction. Whole seedlings (2 g fresh weight) were ground for 2 min with a mortar and pestle on ice in 2 mL extraction buffer (2 mM EGTA, 2 mM EDTA, 100 mM MOPs, pH 7.0) with 1X protease inhibitor cocktail. Extracts were filtered through Miracloth and centrifuged at 10,000 *g* for 10 min to remove cellular debris. A crude microsomal pellet was prepared by centrifuging the supernatant at 100,000 *g* for 1 h at 4°C. The pellet was resuspended in 0.5 ml of resuspension buffer (extraction buffer plus 2% (v/v) Triton X-100) followed by incubation on ice and shaking for 30 min. The supernatant was clarified by further centrifugation at 100,000 *g* for 30 min at 4 °C to remove insoluble debris. Extracted microsomal protein (100 μg per lane) was loaded onto BN-PAGE gels following previously published methods (Atanassov et al., 2009). After electrophoresis, the gel was heated in denaturing buffer (3.3% (w/v) SDS, 65 mM Tris-HCl, pH 6.8) using a microwave at maximum power for 20 s to denature protein complexes and facilitate transfer to PVDF membrane. Protein band detection followed the protein immunoblot analysis protocol described above.

### Spinning-disk confocal microscopy (SDCM)

For live-cell imaging with SDCM, seedlings were grown vertically in the dark for 3 d and images were collected from hypocotyl epidermal cells about 2 mm below the hook. The seedlings were placed in 100 μL of ½-strength MS liquid medium applied onto the slide. A 22 × 40 mm cover glass was positioned on top of the specimen, separated by double-sided tape, for imaging. To examine the cellular localization of YFP-CESA6, imaging was performed using a CSU-X1A1 Yokogawa scanning unit mounted on an Olympus IX83 microscope equipped with a 100X/1.4 NA UPlanSApo oil objective (Olympus) and an Andor iXon Ultra 897BV EMCCD camera (Andor Technology). YFP, mCerulean and mCherry fluorescence were excited with 515-nm, 445-mn and 561-nm laser lines and emission collected through 542/27-nm, 479/40-nm, and 607/36-nm filters, respectively.

For fluorescence recovery after photobleaching (FRAP) experiments, images were collected using a Zeiss Observer Z.1 microscope, equipped with a Yokogawa CSU-X1A1 head and a 100X/1.46 NA PlanApo objective (Zeiss). For the PM-localized CESA6, photobleaching was performed with a Vector scanner (Intelligent Imaging Innovations, Denver, CO) with a 515-nm laser line at 100% power and 1 ms/scan. Time-lapse images were collected at the PM with 10-s intervals for 64 frames, with photobleaching in a small region (44.2 μm^2^) after the 4th frame, and recovery for a total of 5 min. For Golgi-localized CESA6, seedlings were treated with Latrunculin B (6 μM) for 30 min to immobilize the Golgi and time-lapse images were collected at the cortical cytoplasm (about 0.4 μm below the PM) with 5-s intervals for 64 frames. Photobleaching of a small region (7.1 μm^2^) was performed after the 4th frame, and recovery measured for 5 min.

### SDCM image processing and quantification

All image analysis was performed using FIJI/ImageJ (Schindelin et al., 2012). Detailed methods for imaging quantification and analyses were as described previously (Huang et al., 2020; 2021; Zhang et al., 2019; 2021b; Zhang and Staiger, 2022). All imaging experiments were conducted with at least 3 biological repeats.

For analyses of CSC density at the PM, regions of interest (ROIs) without abundant Golgi signals were chosen using the free-hand selection tool. CSCs were detected automatically on 8-bit images using the Find Maxima tool with the same noise threshold for all images. CSC density for each ROI was calculated by dividing the number of particles by the ROI area.

To analyze CSC motility in the plane of the PM, 5-min time-lapse series with 5-s acquisition intervals were collected. Average intensity projections were generated to identify the trajectories of CSCs. Image drift was corrected using the StackReg plugin (Thevenaz et al., 1998). Kymographs were generated, and the speed of CSCs was measured as the reciprocal of the slope of individual particles in the kymographs.

To analyze “long-pause” CSCs, the same 5-min time-lapse series used for CSC speed quantification were used. An ROI of 60 × 60 pixels at the PM was chosen. To create kymographs that cover all of the particle trajectories in the ROI, the Reslice function in FIJI was used and “start at left” was chosen. All vertical lines on kymographs that lasted for 3 min (85 s + 93 s; mean + 3 SD of pause phase in wild type) or longer were counted as “long-pause” CSCs.

To quantify cortical and subcortical YFP-CESA6 containing vesicles or SmaCCs, 1μm z-series stack with 0.2 μm as the step size and 20-s time-lapses with 2-s acquisition intervals were collected. The focal plane at 0.4 μm below the PM was used for cortical SmaCC analysis. To quantify subcortical vesicles, the focal plane from 0.6 μm to 1.0 μm defined as subcortical cytoplasm. Small particles showing motility in time-lapse series were considered to be SmaCCs.

To quantify Golgi-localized YFP-CESA6 fluorescence, 20-s timelapse series with 2-s acquisition intervals used for CSC vesicle quantification were used. Single Golgi with YFP- CESA6 signal and which did not cluster with others were selected for intensity quantification. A square ROI with an area of 3.5 μm^2^ was drawn to include the whole Golgi for analyses. The integrated fluorescence intensity was calculated by subtracting the background fluorescence intensity outside of the cell with the same size of area.

To quantify the ring diameter of Golgi-localized YFP-CESA6, 20-s timelapse series with 2-s acquisition intervals were used. Single Golgi localized YFP-CESA6 which showed typical Golgi shape from top view and did not cluster with others were selected for quantification. A straight line was drawn across the center of Golgi-localized YFP-CESA6 to obtain the plot profile. The distance between two peaks of fluorescence intensity in the plot profile was defined as the ring diameter.

To quantify Golgi-localized YFP-CESA6 and Golgi marker fluorescence intensities in YFP-CESA6 Golgi double marker lines, heterozygous F1 generation seedlings were used. Timelapse series images of 20 s with 2-s acquisition intervals were collected. Single Golgi which did not cluster with others were selected for intensity quantification. The same Golgi colocalized YFP-CESA6 was chosen for Golgi localized CESA6 fluorescence intensity quantification. A square ROI with an area of 3.5 μm^2^ was drawn to include the whole Golgi for analyses. The integrated fluorescence intensity was calculated by subtracting the background fluorescence intensity outside of the cell with the same size of area.

For the FRAP assay of PM-localized CSCs, a smaller ROI (16 μm^2^) within the bleached area was used for analyses. The CSC delivery events during the first 5 min of recovery were manually counted according to the criteria described previously (Li et al., 2016). The particles which exhibited steady linear movement at the PM were considered as new delivery events. The CSC delivery rate was calculated by dividing the number of delivery events by the measured area and time.

For FRAP assay of Golgi-localized YFP-CESA6, an area (4 μm^2^) within the bleached region was used for analyses. To measure the fluorescence intensity, the integrated fluorescence intensity at the selected region at different time points was calculated by subtracting the background fluorescence outside of the cell with the same size of area. Based on the slope of fluorescence recovery curve, the mean recovery rate was calculated.

### Correlation and principal component analyses

Pairwise correlation analyses between cellulose content, hypocotyl length, and various CSC trafficking parameters were conducted using JMP 11 software. Each parameter was normalized to wild-type YFP-CESA6 and the corresponding relative parameter data was used for the correlation analysis. Bivariate fit/ANOVA was performed for all data points for each parameter. Principal component analysis was conducted using JMP 11 software.

### Homology modeling

Homology modeling of the full-length AtCESA6 structure, using PttCESA8 as the template (Purushotham et al., 2020), was performed by using the SWISS-MODEL server as described previously (Huang et al., 2021). Figures were prepared using PyMOL v1.7.6.7 software.

### Accession numbers

Amino acid sequence for AtCESA6 can be found in the UniProt Knowledgebase under accession number Q94JQ6. The gene accession number for *AtCESA6* in TAIR (The Arabidopsis Information Resource) is AT5G64740.

## Supplemental data

The following materials are available in the online version of this article.

**Supplemental Table S1.** Catalog of mutations and complementation lines for YFP-CESA6.

**Supplemental Table S2.** Summary of predicted effects of CESA6 mutations on cellulose synthesis.

**Supplemental Table S3.** Summary of effects of CESA6 mutations on growth, cellulose content, and vesicle trafficking/dynamics.

**Supplemental Table S4.** Primers used to generate YFP-CESA6 site directed mutagenesis constructs and perform RT-qPCR.

**Supplemental Figure S1.** The original *es20r* EMS mutants generally exhibit reduced growth and cellulose content in dark grown hypocotyls.

**Supplemental Figure S2.** Mutations in key motifs of the CESA6 catalytic domain affect hypocotyl growth.

**Supplemental Figure S3**. Expression levels of YFP-CESA6 in wild type and 18 mutated CESA6 transgenic lines are comparable.

**Supplemental Figure S4.** Distribution of YFP-CESA6 particle speeds from transgenic plants expressing mutated YFP-CESA6.

**Supplemental Figure S5.** Analysis of a second independent transgenic line shows similar CSC dynamics and Golgi fluorescence intensity phenotypes when compared with the original transgenic line.

**Supplemental Figure S6.** Abundance of the Golgi markers mCerulean-MEMB12 and mCherry-SYP32 are not affected by mutations in the DDG and QXXRW motifs.

**Supplemental Figure S7.** BFA treatment does not cause abnormal accumulation of cortical SmaCCs in YFP-CESA6 mutant lines.

**Supplemental Figure S8.** Crystalline cellulose content positively correlates with hypocotyl length and negatively correlates with cortical SmaCC density and Golgi fluorescence intensity

**Supplemental Figure S9.** Principal component analysis of cellulose biosynthesis and CSC trafficking parameters.

**Supplemental Figure S10.** Group II homologous mutations are close to the homodimer interface formed by β-strand 6 (β6) and α-helix 9 (α9) in AtCESA3 catalytic domain structure.

## Acknowledgments

This work is dedicated to the memory of Dr. Chunhua Zhang, a brilliant scientist, dear friend, and valued colleague who passed away on May 15, 2021. We are grateful to Daniel Szymanski (Purdue University) for sharing the spinning disc confocal microscope for FRAP studies and to Ruthie Arieti (UNC Chapel Hill) for guidance on correlation and PCA analyses. We are grateful to Yuxian Zhu and Xingpeng Wen (Wuhan University) for guidance on microsome extraction and BN-PAGE assays. This research was supported by the Purdue University Provost Start-up Fund to C.Z., and a US National Science Foundation grant (MCB-2025437) to C.Z. and C.J.S.

## Author contributions

L.H., W.Z., X.L., C.Z., and C.J.S. conceived the experiments and designed the research; L.H. and W.Z. performed experiments and analyzed data; L.H., W.Z., and C.J.S. wrote the article.

