## Supplemental Data for "Point mutations in the catalytic domain disrupt cellulose synthase (CESA6) vesicle trafficking and protein dynamics"

**Supplemental Table S1.** Catalog of mutations and complementation lines for YFP-CESA6.

| Motif/Domain | Mutations |
|---|---|
| DDG | S394F#, D395N§, D396N# |
| DXD | D562N§, D564N§ |
| TED | T783I#, D785N§ |
| QXXRW | Q823E§, R826A§, W827A§ |
| IF3 | E929K#, G935E# |
| ES20 | L365F§, S360N#, P595S#, D602N#, D605N#, G632D# |

§ Site-directed mutagenesis of putative ES20-binding pocket based on homology model of CESA6 threaded onto bacterial CESA (Huang et al., 2020)

### Denotes amino acid change/mutation discovered in *es20r* screen and recreated in YFP-CESA6 complementation line (Huang et al., 2020)

**Supplemental Table S2.** Summary of predicted effects of CESA6 mutations on cellulose synthesis

| Mutations | Motifs | Predicted effect‡ |
|---|---|---|
| **S360N#** | | Not fully understood; the mutation may affect UDP-Glc binding to acceptor |
| **L365F§** | | Not fully understood; the mutation may affect UDP-Glc binding to acceptor |
| **S394F#** | **DDG** | Affects coordination of UDP and hence occludes UDP-Glc binding to acceptor |
| **D395N§** | **DDG** | Affects coordination of UDP and hence occludes UDP-Glc binding to acceptor |
| **D396N#** | **DDG** | D396 contacts the uracil moiety of UDP through hydrogen bond; the mutation likely affects coordination of UDP and hence occludes UDP-Glc binding to acceptor |
| **D562N§** | **DXD** | D562 may contact with diphosphate moiety of UDP through hydrogen bond; the mutation likely affects coordination of UDP and hence occludes UDP-Glc binding to acceptor |
| **D564N§** | **DXD** | D564 may contact with phosphate moiety of UDP through hydrogen bond; the mutation affects coordination of UDP and hence occludes UDP-Glc binding to acceptor |
| **P595S#** | | Unclear |
| **D602N#** | | Unclear |
| **D605N#** | | Unclear |
| **G632D#** | | Unclear |
| **T783I#** | **TED** | TED is part of finger helix and facilitates the glucose chain translocation. T783 forms hydrogen bond with terminal glucose's C2 hydroxyl group and the mutation may affect the interaction with glucan chain's terminal glucose and hence disturbs the driving force which pushes the elongated glucan chain in transmembrane channel |
| **D785N§** | **TED** | TED is part of finger helix and facilitates the glucose chain translocation. D785 forms hydrogen bond with terminal glucose's C4 hydroxyl group and facilitates the deprotonation of the C4 hydroxyl. The mutation may affect the interaction with glucan chain's terminal glucose and hence disturbs the driving force which pushes the elongated glucan chain in transmembrane channel |
| **Q823E§** | **QXXRW** | Q823 forms hydrogen bond with the neighbor residue; mutation may affect interaction of glucose with glucan chain at the acceptor-binding site and hence interfere with glucan chain elongation |
| **R826A§** | **QXXRW** | R826 likely coordinates the diphosphate of UDP; mutation may affect interaction of glucose with glucan chain at the acceptor-binding site and hence interfere with glucan chain elongation |
| **W827A§** | **QXXRW** | W827 forms van der Waals interaction with the penultimate glucose; the mutation may affect interaction of glucose with glucan chain at the acceptor-binding site and hence interfere with glucan chain elongation |

| | | |
|---|---|---|
| **E929K#** | **IF3** | Mutation may affect the channel across membrane and hence interfere with glucan chain translocation |
| **G935E#** | **IF3** | Mutation may affect the channel across membrane and hence interfere with glucan chain translocation |

‡ Summarized from the following references: Morgan et al., 2013, McNamara et al., 2015, Morgan et al., 2016 and Purushotham et al., 2020.

§ Site-directed mutagenesis of putative ES20-binding pocket based on homology model of CESA6 threaded onto bacterial CESA (Huang et al., 2020)

### Denotes amino acid change/mutation discovered in *es20r* screen and recreated in YFP-CESA6 complementation line (Huang et al., 2020)

**Supplemental Table S3.** Summary of effects of CESA6 mutations on growth, cellulose content, and vesicle trafficking/dynamics

| Group | Mutation | Motif | Hypocotyl growth (*es20r* lines) | Cellulose content (*es20r* lines) | Hypocotyl growth (YFP-CESA6 lines) | Cellulose content (YFP-CESA6 lines) | CSC PM speed | Long-pause CSC | Golgi fluor. intensity | Golgi fluor. recovery | Golgi ring diameter | Cortical SmaCC density | CSC delivery rate | CSC PM density |
|---|---|---|---|---|---|---|---|---|---|---|---|---|---|---|
| | S360N[#] | | – (-5.1%) | ↓ (-8.1%) | ↓ (-39.1%) | ↓ (-26.2%) | – (-6.8%) | – (0%) | – (+10.9%) | | – (+1.1%) | – (+5.0%) | – (-9.0%) | – (-13.2%) |
| | L365F[§] | | | | – (-2.3%) | ↓ (-11.4%) | – (+4.4%) | – (+15.4%) | ↑ (+126.4%) | ↑ (+88.8%) | – (+12.8%) | ↑(+42.5%) | ↓ (-52.0%) | ↓ (-49.6%) |
| | S394F[#] | DDG | – (0.0%) | ↓ (-11.1%) | ↓ (-35.5%) | ↓ (-17.9%) | – (+1.9%) | – (0%) | ↑ (+136.1%) | ↑ (+166.5%) | – (+7.1%) | – (-3.1%) | ↓ (-45.6%) | ↓ (-61.8%) |
| | D395N[§] | DDG | | | ↓ (-22.0%) | ↓ (-15.9%) | – (+3.2%) | – (-28.6%) | ↑ (+83.8%) | ↑ (+66.1%) | – (+13.3%) | – (+12.5%) | ↓ (-58.9%) | ↓ (-44.0%) |
| Group I | D396N[#] | DDG | ↓ (-40.1%) | ↓ (-16.9%) | ↓ (-25.8%) | ↓ (-12.6%) | – (+6.7%) | – (-28.6%) | ↑ (+37.4%) | – (+49.8%) | – (+10.5%) | – (-17.6%) | ↓ (-71.1%) | – (-9.6%) |
| | D602N[#] | | ↓ (-25.4%) | – (-5.5%) | ↓ (-55.2%) | ↓ (-25.5%) | ↓ (-16.0%) | ↑ (+1899.7%) | ↑(+237.5%) | ↑ (+45.0%) | – (+4.3%) | ↑(+48.3%) | ↓ (-61.3%) | ↓ (-71.7%) |
| | D605N[#] | | ↓ (-17.5%) | ↓ (-10.9%) | ↓ (-67.3%) | ↓ (-27.9%) | ↓ (-46.6%) | ↑ (+1187.5%) | ↑ (+68.9%) | | – (+0.8%) | – (-17.8%) | ↓ (-58.5%) | – (-18.6%) |
| | E929K[#] | IF3 | – (0.0%) | ↓ (-10.4%) | – (-8.3%) | ↓ (-14.9%) | – (+6.3%) | – (+20.0%) | – (+18.3%) | | –(-3.5%) | – (+6.8%) | – (-7.6%) | ↓ (-16.7%) |
| | G935E[#] | IF3 | – (0.0%) | – (-6.7%) | – (-12.1%) | ↓ (-10.3%) | ↓ (-23.8%) | ↑ (+2000.4%) | ↑ (+101.6%) | | –(-2.2%) | – (-10.7%) | ↓ (-65.6%) | ↓ (-58.5%) |
| | D562N[§] | DXD | | | ↓ (-31.5%) | ↓ (-11.7%) | ↓ (-20.0%) | ↑ (+1214.0%) | – (-7.1%) | | – (+11.3%) | ↓ (-30.1%) | ↓ (-60.9%) | – (+2.0%) |
| Group II | D564N[§] | DXD | | | ↓ (-24.7%) | ↓ (-13.2%) | ↓ (-41.8%) | ↑ (+1514.1%) | ↓(-48.2%) | –(-32.6%) | –(-1.5%) | ↓ (-57.7%) | ↓ (-56.3%) | ↑ (+35.6%) |
| | P595S[#] | | – (-5.4%) | ↓ (-15.9%) | ↓ (-15.9%) | ↓ (-7.9%) | ↓ (-21.7%) | ↑ (+1977.0%) | – (-12.8%) | – (+6.4%) | – (+0.8%) | ↓ (-41.4%) | ↓ (-55.0%) | – (+1.0%) |

| | | | | | | | | | | | | | | |
|---|---|---|---|---|---|---|---|---|---|---|---|---|---|---|
| Group II | G632D[#] | | ↓ (-22.0%) | ↓ (-10.7%) | ↓ (-46.0%) | ↓ (-10.9%) | ↓ (-50.8%) | ↑ (+950.3%) | – (-10.6%) | | – (+4.7%) | ↓ (-31.2%) | ↓ (-39.4%) | – (+1.0%) |
| | T783I[#] | TED | – (0.0%) | ↓ (-16.8%) | ↓ (-18.2%) | ↓ (-10.7%) | ↓ (-12.9%) | ↑ (+960.3%) | – (-4.6%) | | – (+7.9%) | – (-27.2%) | ↓ (-52.7%) | – (-2.1%) |
| | D785N[§] | TED | | | ↓ (-49.9%) | ↓ (-13.4%) | ↓ (-41.8%) | ↑ (+1420.3%) | – (+10.5%) | | – (+13.4%) | ↓(-48.7%) | ↓ (-64.1%) | – (-5.9%) |
| | Q823E[§] | QXX RW | | | ↓ (-32.3%) | ↓ (-12.0%) | ↓ (-24.5%) | ↑ (+1313.5%) | ↓ (-52.0%) | ↓ (-25.4%) | ↓ (-14.7%) | ↓ (-54.4%) | ↓ (-43.8%) | ↑ (+25.7%) |
| | R826A[§] | QXX RW | | | ↓ (-26.1%) | ↓ (-10.1%) | ↓ (-49.9%) | ↑ (+1660.5%) | ↓ (-48.2%) | ↓ (-25.1%) | ↓ (-18.8%) | ↓ (-45.4%) | ↓ (-50.0%) | – (+11.9%) |
| | W827A[§] | QXX RW | | | ↓ (-27.4%) | ↓ (-14.9%) | ↓ (-31.6%) | ↑(+1346.5%) | ↓ (-21.3%) | –(-20.6%) | –(-1.6%) | – (-16.2%) | ↓ (-64.4%) | – (-19.8%) |
| | L286F | | ↓ (-26.4%) | ↓ (-24.1%) | | | | | | | | | | |
| | G780S | | – (+15.1%) | ↓ (-7.4%) | | | | | | | | | | |
| | G811E | | ↓ (-51.9%) | ↓ (-11.6%) | | | | | | | | | | |
| | S818T | | – (+9.3%) | – (-0.9%) | | | | | | | | | | |
| | L829F | | ↓ (-19.3%) | ↓ (-13.6%) | | | | | | | | | | |
| | L1029F | | – (+12.3%) | ↓ (-7.1%) | | | | | | | | | | |

[§] Site-directed mutagenesis of putative ES20-binding pocket based on the homology model of CESA6 threaded onto bacterial CESA (Huang et al., 2020)

[#] Denotes amino acid change/mutation discovered in *es20r* screen and recreated in YFP-CESA6 complementation line (Huang et al., 2020)

"–": no difference with control; "↓": reduced; "↑": increased; percentage are relative to Col-0 or wild-type YFP-CESA6 in *prc1-1*

**Supplemental Table S4.** Primers used to generate YFP-CESA6 site directed mutagenesis constructs and perform qRT-PCR

| Primer name | Primer sequence (5'-3') |
|---|---|
| CESA6G935E-F | GGGTCATTGAAGGTGTTTCTGC |
| CESA6G935E-R | AAAACTGTTCGTTTCTCCACCAA |
| CESA6P595S-F | GTTCAGTTCTCTCAAAGGTTCGA |
| CESA6P595S-R | ATAACAGATTTTCTTTCCTGACT |
| CESA6D605N-F | ATTGATAGGCACAATCGATACTCAA |
| CESA6D605N-R | CCCATCGAACCTTTGAGGGAACTG |
| CESA6G632D-F | GTCGATACAGGTTGTGTTTTCA |
| CESA6G632D-R | GTATATAGGCCCTTGTAGCCCAT |
| CESA6S394F-F | GTTACGTATTTGATGATGGTGCT |
| CESA6S394F-R | AAGCAACCTTATCGACAGGATA |
| CESA6D602N-F | TGGGATTAATAGGCACGATCG |
| CESA6D602N-R | TCGAACCTTTGAGGGAACTG |
| CESA6S360N-F | GATGTATTTGTTAATACAGTGGATCC |
| CESA6S360N-R | CACAGGGGATAGTCCCGAC |
| CESA6D396N-F | GTATCTGATAATGGTGCTGCTA |
| CESA6D396N-R | GTAACAAGCAACCTTATCGAC |
| CESA6T783I-F | GGTTCTGTTATCGAAGATATTCT |
| CESA6T783I-R | ATAGATCCACCCAATCTGAGA |
| CESA6E929K-F | GGAGAAACAAACAGTTTTGGGT |
| CESA6E929K-R | ACCAATCATCGATCCCAACTTT |
| CESA6L365F-F | GGATCCATTTAAAGAGCCTCCG |
| CESA6L365F-R | ACTGTACTAACAAATACATCCAC |
| CESA6D395N-F | ACGTATCTAATGATGGTGCT |
| CESA6D395N-R | AACAAGCAACCTTATCGACA |
| CESA6D562N-F | CTGAATGTCAATTGTGATCAC |
| CESA6D562N-R | AAGGTAAGGAGCATTTGATA |
| CESA6D564N-F | GTCGATTGTAATCACTACATCA |
| CESA6D564N-R | ATTCAGAAGGTAAGGAGCATTT |
| CESA6D785N-F | GTTACCGAAAATATTCTTACGGG |
| CESA6D785N-R | AGAACCATAGATCCACCCAATCTG |
| CESA6Q823E-F | CGTCTCCATGAAGTTCTTCGAT |
| CESA6Q823E-R | ATCCGAAAGATTGATTGGAGCT |
| CESA6R826A-F | AAGTTCTTGCATGGGCGCTTGG |
| CESA6R826A-R | GATGGAGACGATCCGAAAGATT |
| CESA6W827A-F | TTCTTCGAGCGGCGCTTGGGTCG |
| CESA6W827A-R | CTTGATGGAGACGATCCGAAAGA |
| CESA6qPCR-F | ACTACCTGAGCTACCAGTCCG |
| CESA6qPCR-R | CCATCAACAGTCAATTCGATCTC |
| ACTIN1qPCR-F | GGCGATGAAGCTCAATCCAAA |
| ACTIN1qPCR-R | GGTCACGACCAGCAAGATCAA |

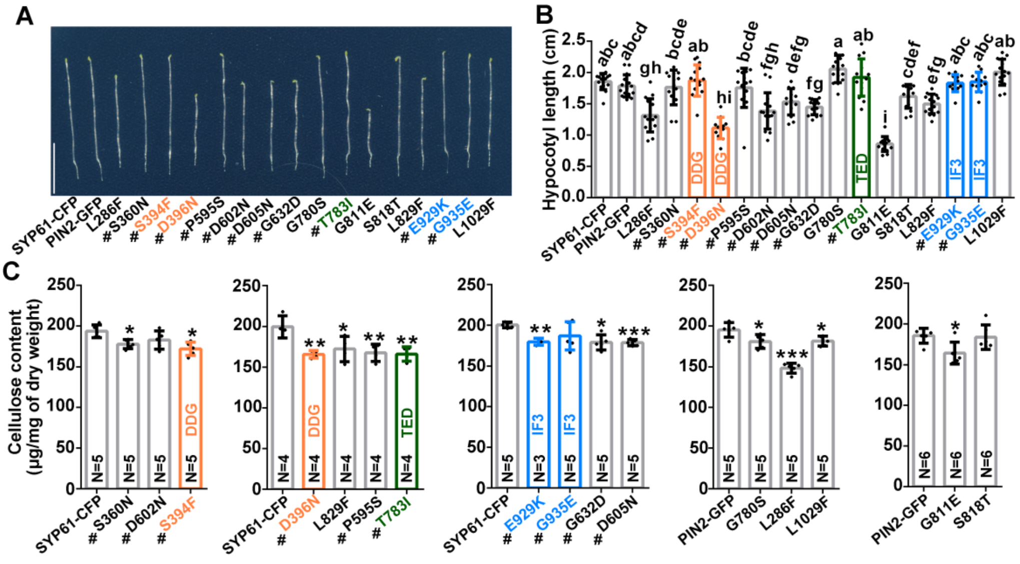

**Figure S1.** The original *es20r* EMS mutants generally exhibit reduced growth and cellulose content in dark grown hypocotyls. (Supports Figure 1).

**(A)** Representative 7-d-old dark grown seedlings of EMS-induced mutants that exhibit reduced sensitivity to ES20. Scale bar: 1 cm. SYP61-CFP and PIN2-GFP refer to the EMS-mutagenized lines used to screen for ES20 resistance phenotypes. # Denotes amino acid change/mutation discovered in *es20r* screen and recreated in YFP-CESA6 complementation line (Huang et al., 2020). **(B)** Quantification of hypocotyl length for seedlings described in A. Mutations in the conserved catalytic core motifs DDG and TED, as well as Interfacial helix3 (IF3) motifs are highlighted with corresponding colors. Data represent mean ± SD ($n \geq 11$ seedlings per genotype). Statistically significant differences were determined using one-way analysis of variance (ANOVA) followed by Tukey's multiple comparisons test. Different letters indicate significant differences between groups ($p < 0.05$). **(C)** Crystalline cellulose content in hypocotyl cell walls of 7-day-old dark-grown EMS-induced *cesa6* mutants. Data represent mean ± SD. Dots on each bar represent an independent biological replicate prepared from seedlings of the same genotype. Each biological replicate value was obtained by calculating the mean of at least three technical replicates. * indicates $p < 0.05$, and *** indicates $p < 0.001$ by one-way ANOVA followed by Dunnett's multiple comparisons test compared with wild-type SYP61-CFP or PIN2-GFP lines.

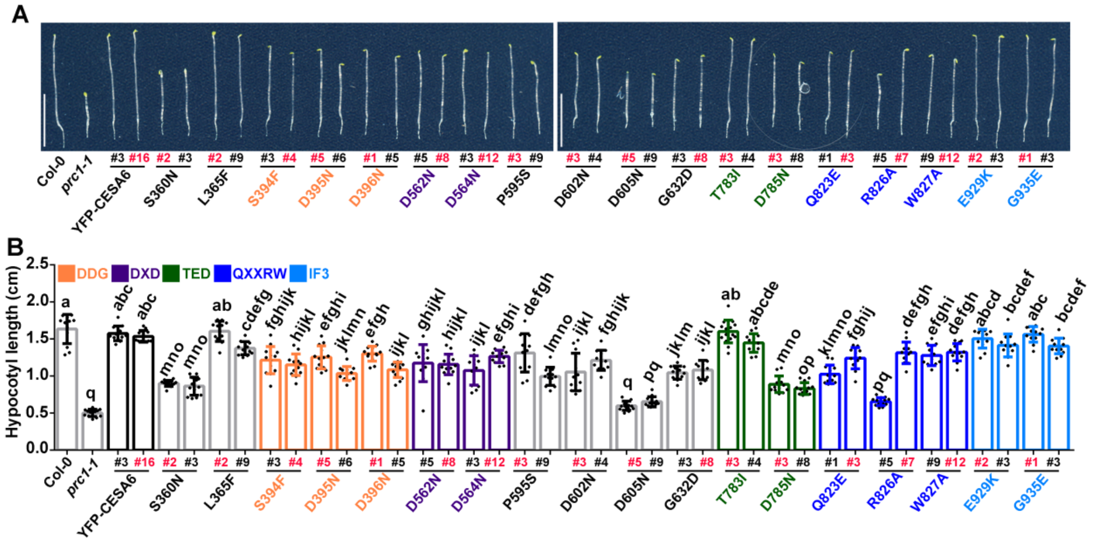


**Figure S2.** Mutations in key motifs of the CESA6 catalytic domain affect hypocotyl growth. (Supports Figure 1)

**(A)** Representative 3-d-old dark grown seedlings of two independent transgenic lines expressing wild-type or mutated YFP-CESA6. Scale bars: 1 cm. Numbers in red represent the transgenic mutant line used for cellulose quantification and live cell imaging analysis in the main text. **(B)** Quantification of hypocotyl length for 3-d-old dark grown seedlings as described in A. Data represent mean ± SD ($n \geq 11$ seedlings per genotype). Statistically significant differences were determined using one-way analysis of variance (ANOVA) followed by Tukey's multiple comparisons test. Different letters indicate significant differences between groups ($p < 0.05$).

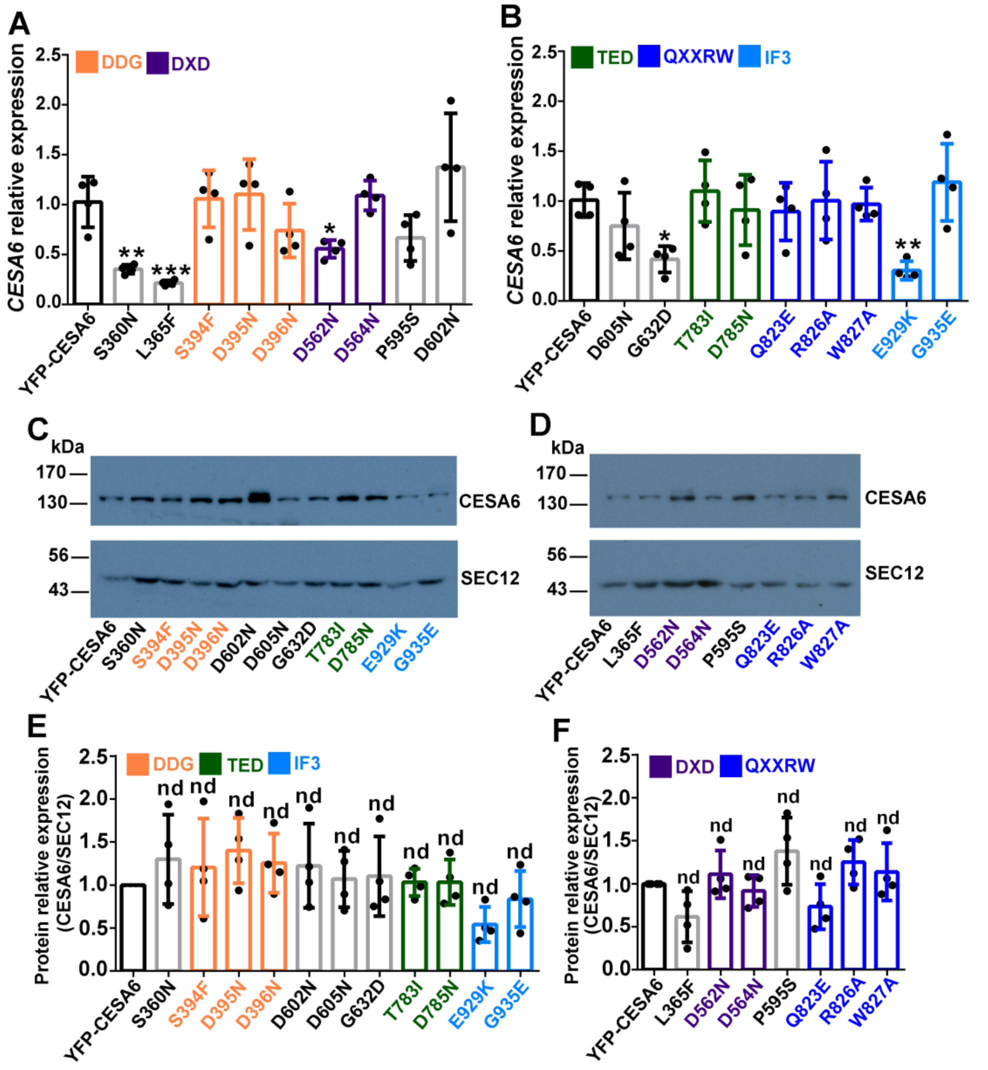


**Figure S3.** Expression levels of YFP-CESA6 in wild type and 18 mutated CESA6 transgenic lines are comparable. (Supports Figure 1)

(**A** and **B**) Quantification of *CESA6* transcript level in wild-type and mutated YFP-CESA6 transgenic lines using qPCR. Data represent mean ± SD (n = 4 biological replicates). * indicates $p < 0.05$, ** indicates $p < 0.01$ and *** indicates $p < 0.001$ by one-way ANOVA followed by Dunnett's multiple comparisons test compared with wild-type YFP-CESA6. (**C** and **D**) Representative protein immunoblots show the abundance of YFP-CESA6 and SEC12 in total protein extracts isolated from seedlings of wild type and 18 different transgenic lines expressing mutated YFP-CESA6 in *prc1-1*. The membrane was cut between 95-kDa and 72-kDa markers and probed separately. The upper membrane was probed with anti-GFP antibody for YFP-CESA6 detection and the lower membrane was probed with anti-SEC12 antibody as a loading control. (**E** and **F**) Quantification of the relative abundance of YFP-CESA6 in wild type and 18 different transgenic lines using immunoblot analysis described in C and D. Integrated intensity of each immunoblot band was measured with ImageJ and background intensity was subtracted from each measurement. For each sample, the abundance of YFP-CESA6 was normalized to SEC12 level. Normalized YFP-CESA6 abundance in each sample was then compared with wild-type YFP-CESA6 sample. Data represent mean ± SD (n = 4 biological replicates). nd indicates no significant difference. Significant difference was analyzed by one-way ANOVA followed by Dunnett's multiple comparisons test compared with wild-type YFP-CESA6.

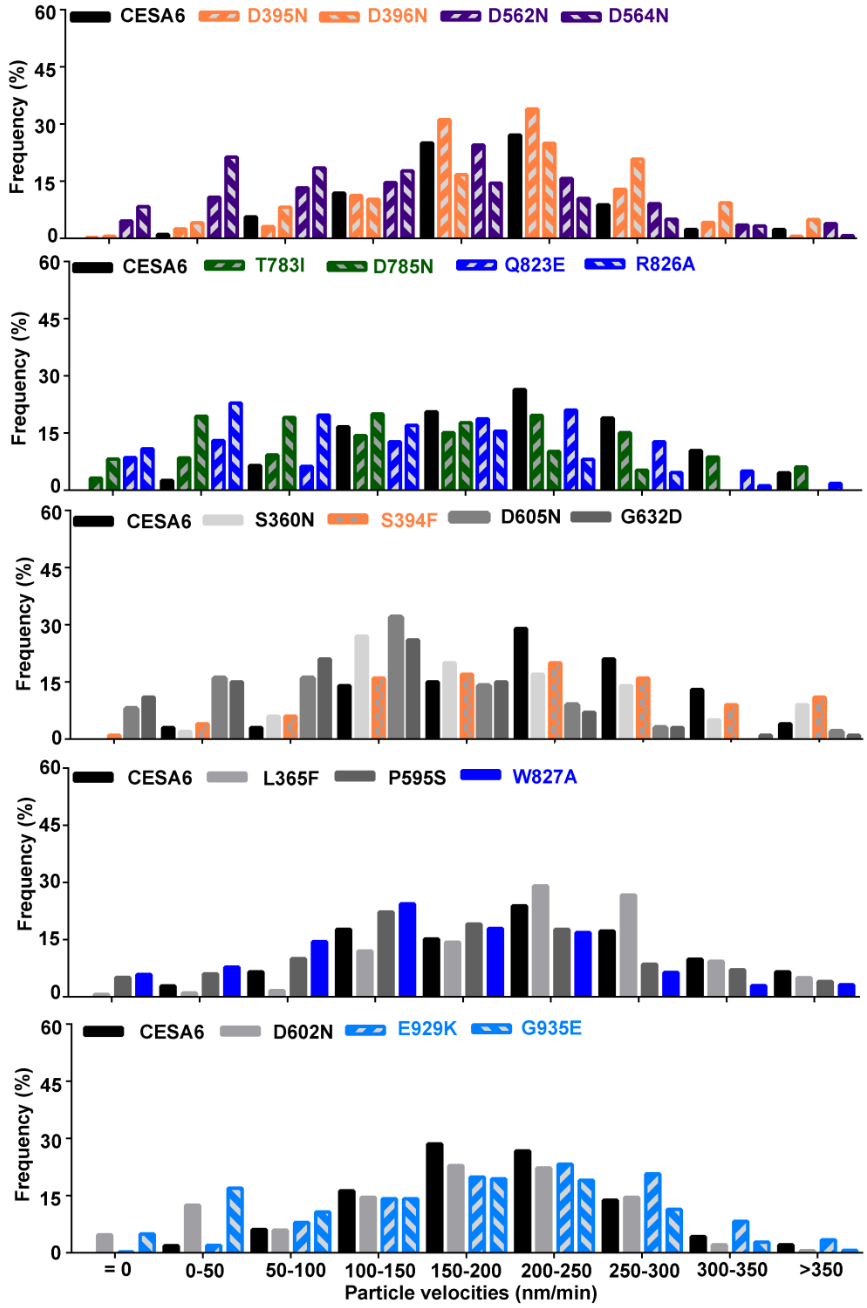


**Figure S4.** Distribution of YFP-CESA6 particle speeds from transgenic plants expressing mutated YFP-CESA6. (Supports Figure 2)

Mutations in the DXD, TED, and QXXRW motifs left skewed the frequency distribution and caused appearance of a population of static CSC particles at the PM.

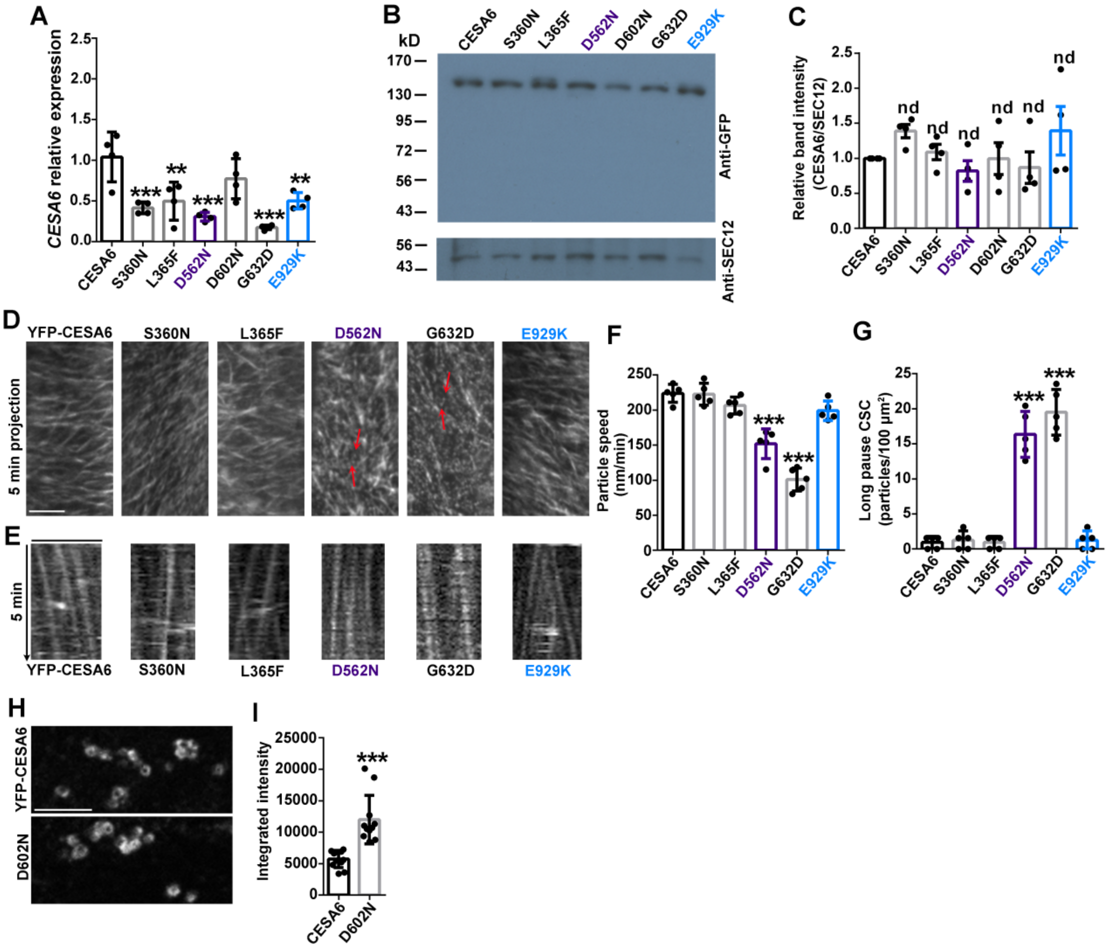


**Figure S5.** Analysis of a second independent transgenic line shows similar CSC dynamics and Golgi fluorescence intensity phenotypes when compared with the original transgenic line. (Supports Figure 2-4)

(**A**) Quantification of *CESA6* transcript levels in wild-type and mutated YFP-CESA6 transgenic lines using qRT-PCR. Data represent mean ± SD (n = 4 biological replicates). ** indicates p < 0.01 and *** indicates p < 0.001 by one-way ANOVA followed by Dunnett's multiple comparisons test compared with wild-type YFP-CESA6. (**B**) Representative protein immunoblots show the abundance of YFP-CESA6 and SEC12 in total protein extracts isolated from seedlings of wild type and mutated YFP-CESA6 transgenic lines. (**C**) Quantification of the relative abundance of YFP-CESA6 in wild type and different transgenic lines using immunoblot analysis described in B. Data represent mean ± SD (n = 4 biological replicates). Significant differences were analyzed by one-way ANOVA followed by Dunnett's multiple comparisons test compared with wild-type YFP-CESA6. nd indicates no significant difference. **(D)** Representative time projections prepared with average intensity images from a time-lapse series of YFP-CESA6 particles at the PM in hypocotyl epidermal cells of transgenic plants expressing mutated CESA6. Red arrows highlight static CSC particles present in some mutants. Scale bar: 5 µm. **(E)** Representative kymographs of the trajectories of YFP-CESA6 particles from seedlings as described in D. Scale bar: 5 µm. **(F)** Quantification of YFP-

CESA6 particle speed. Data represent mean ± SD (n = 5 seedlings per sample, more than 50 CSC trajectories were analyzed for each seedling). *** indicates $p < 0.001$ by one-way ANOVA followed by Dunnett's multiple comparisons test compared with wild-type YFP-CESA6 complementation line. (**G**) Quantification of the density of long-pause CSCs for mutated CESA6 transgenic lines. Data represent mean ± SD (n = 5 seedlings per sample, 2 cells from each seedling were chosen for analysis). *** indicates $p < 0.001$ by one-way ANOVA followed by Dunnett's multiple comparisons test compared with wild-type YFP-CESA6 complementation line. (**H**) Representative images of YFP-CESA6 at the Golgi in etiolated hypocotyl epidermal cells of transgenic plants expressing wild type CESA6 or CESA6-D602N. Scale bar: 5 μm. **(I)** Quantification of the fluorescence intensity of YFP-CESA6 at Golgi as described in H. Data represent mean ± SD (n = 12 seedlings, 5 Golgi from each seedling were chosen for analysis). *** indicates $p < 0.001$ by Student's *t*-test compared with wild-type YFP-CESA6 complementation line.

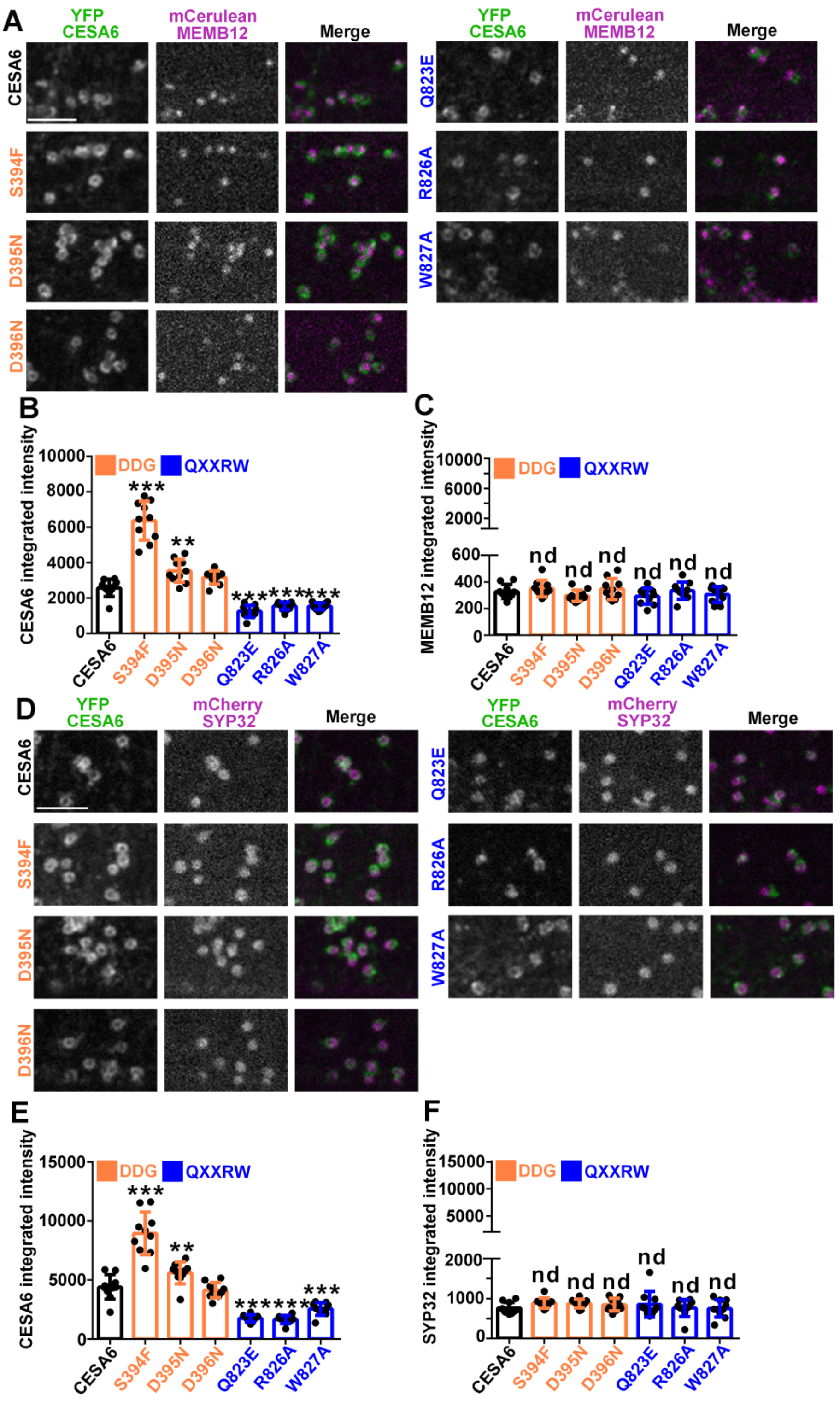


**Figure S6.** Abundance of the Golgi markers mCerulean-MEMB12 and mCherry-SYP32 are not affected by mutations in the DDG and QXXRW motifs. (Supports Figure 4)

**(A)** Representative colocalization images of mCerulean-MEMB12 and YFP-CESA6 in the Golgi of etiolated hypocotyl epidermal cells. Scale bar: 5 µm. **(B and C)** Fluorescence intensity of YFP-CESA6 and mCerulean-MEMB12 in the Golgi, respectively, as described in A. Data represent mean ± SD (n = 10 seedlings, 5 Golgi from each seedling were chose for analysis). nd represents no significant difference, ** indicates $p < 0.01$ and *** indicates $p < 0.001$ by one-way ANOVA followed by Dunnett's multiple comparisons test compared with wild-type YFP-CESA6. **(D)** Representative colocalization images of mCherry-SYP32 and YFP-CESA6 in the Golgi in hypocotyl epidermal cells. Scale bar: 5 µm. **(E** and **F)** Fluorescence intensity of YFP-CESA6 and mCherry-MEMB12 in the Golgi, respectively, as described in D. Data represent mean ± SD (n = 10 seedlings, 5 Golgi from each seedling were chose for analysis). nd represents no significant difference, ** indicates $p < 0.01$ and *** indicates $p < 0.001$ by one-way ANOVA followed by Dunnett's multiple comparisons test compared with wild-type YFP-CESA6.

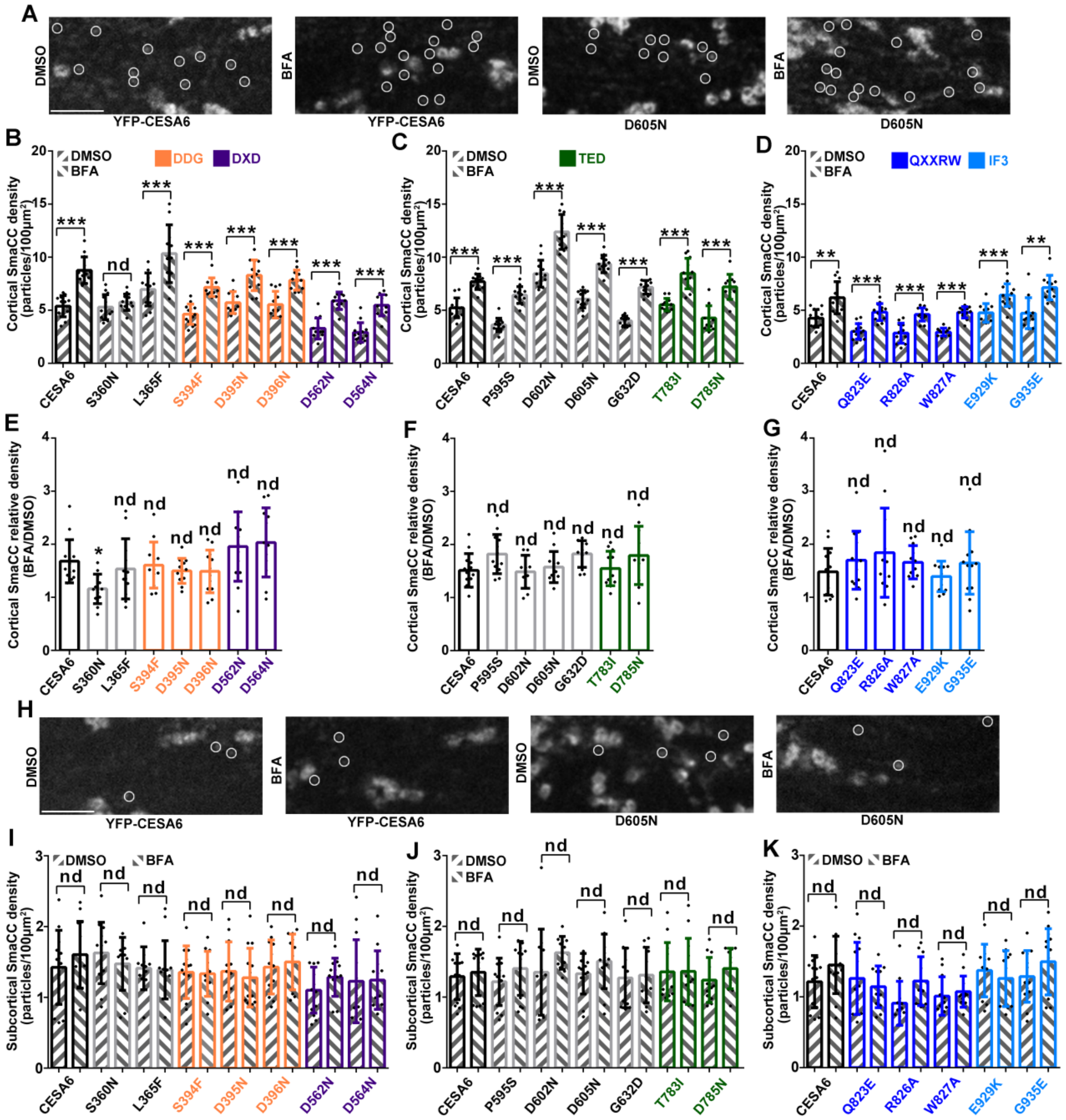


**Figure S7.** BFA treatment does not cause abnormal accumulation of cortical SmaCCs in YFP-CESA6 mutant lines. (Supports Figures 5 and 6)

**(A)** Representative images of SmaCCs in the cortical region of etiolated hypocotyl epidermal cells of transgenic seedlings expressing wild-type or mutated YFP-CESA6 transgenic lines treated with DMSO (0.1%) or BFA (50 µM) for 2 h. White circles indicate designated SmaCCs. Scale bar: 5 µm. **(B–D)** Quantification of cortical SmaCC density as shown in A. Data represent mean ± SD (n ≥ 12 cells from 12 seedlings). nd represents no significant difference, ** indicates $p < 0.01$ and *** indicates $p < 0.001$ by Student's *t*-test. (**E–G**) Quantification of relative cortical SmaCC density of BFA treatment divided by DMSO treatment from wild-type and mutated YFP-CESA6 transgenic lines. nd represents no significant difference. Statistically significant differences were determined using by one-way ANOVA followed by Dunnett's multiple comparisons test compared with wild-type YFP-CESA6. (**H**) Representative images of SmaCCs in the subcortical region of hypocotyl epidermal cells of transgenic seedlings expressing wild-type or mutated YFP-CESA6 transgenic lines treated with DMSO (0.1%) or BFA (50 μM) for 2 h. White circles indicate designated SmaCCs. Scale bar: 5 μm. **(I–K)** Quantification of subcortical SmaCC density as shown in H. Data represent mean ± SD (n ≥ 12 cells from 12 seedlings). nd represents no significant difference by Student's *t*-test.

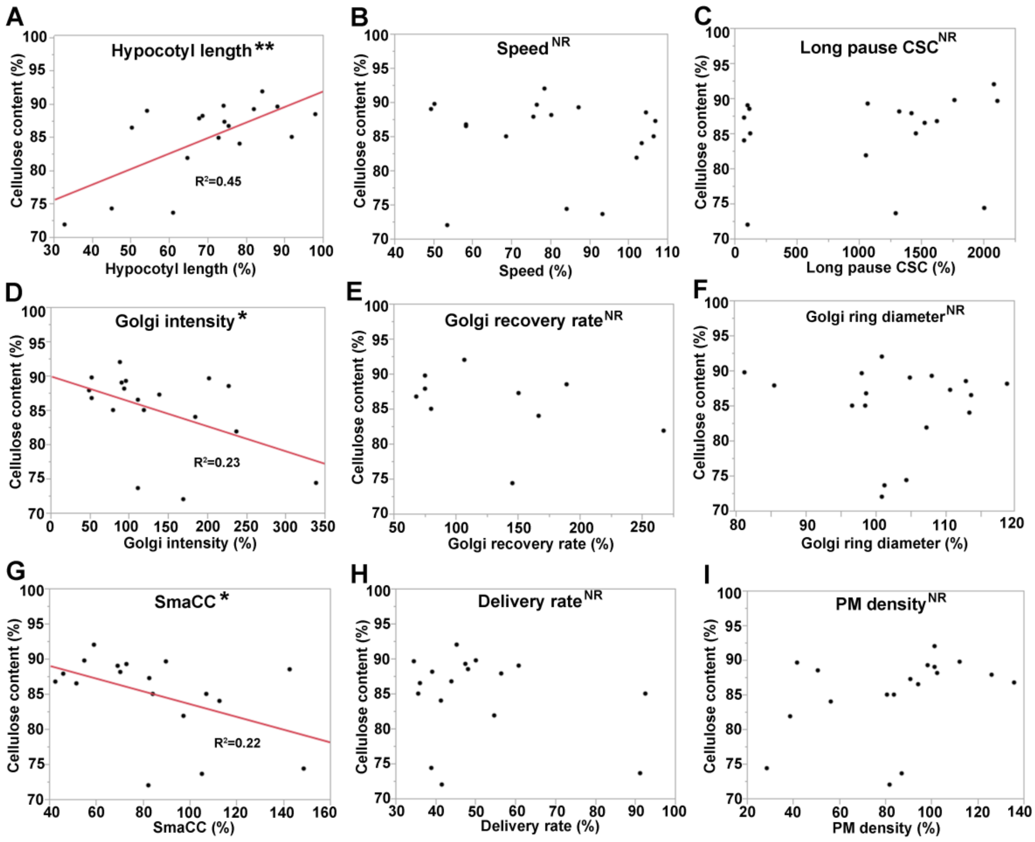


**Figure S8.** Crystalline cellulose content positively correlates with hypocotyl length and negatively correlates with cortical SmaCC density and Golgi fluorescence intensity. **(A–I)** Pairwise correlation analysis between crystalline cellulose content and hypocotyl length or various CSC trafficking parameters was conducted using JMP 11 software. Crystalline cellulose content was plotted as a function of hypocotyl length **(A)**, CSC speed **(B)**, Long-pause CSC abundance **(C)**, Golgi fluorescence intensity **(D)**, Golgi fluorescence recovery rate **(E)**, Golgi ring diameter **(F)**, cortical SmaCC density **(G)**, CSC delivery rate **(H)** and CSC PM density **(I)**. Corresponding parameters in **(A–I)** were collected from 18 YFP-CESA6 mutated complementation lines and Golgi fluorescence recovery rate in **(E)** was obtained from 10 YFP-CESA6 mutated complementation lines. Each parameter was normalized to wild-type YFP-CESA6 and the corresponding relative values were used for the correlation analysis. NR, no predictive relationship; *, $P \leq 0.05$, **, $P \leq 0.01$, Bivariate fit/ANOVA for all data points for each parameter.

A

**Eigenvalues**

| Number | Eigenvalue | Percent | 20 40 60 80 | Cum Percent |
|---|---|---|---|---|
| 1 | 3.7843 | 42.047 | | 42.047 |
| 2 | 2.0301 | 22.557 | | 64.604 |
| 3 | 1.5298 | 16.998 | | 81.602 |
| 4 | 0.9524 | 10.583 | | 92.184 |
| 5 | 0.2773 | 3.082 | | 95.266 |
| 6 | 0.1796 | 1.995 | | 97.262 |
| 7 | 0.1351 | 1.501 | | 98.763 |
| 8 | 0.0732 | 0.813 | | 99.576 |
| 9 | 0.0381 | 0.424 | | 100.000 |

C

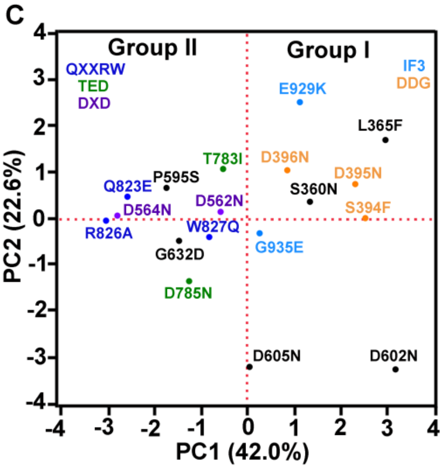


B

**Eigenvectors**

| | Prin1 | Prin2 | Prin3 | Prin4 | Prin5 | Prin6 | Prin7 | Prin8 | Prin9 |
|---|---|---|---|---|---|---|---|---|---|
| Cellulose content | -0.25110 | 0.41216 | 0.46785 | 0.08979 | 0.26306 | -0.41870 | 0.24560 | 0.42098 | -0.24098 |
| Hypocotyl length | 0.01990 | 0.60437 | 0.26246 | 0.34595 | 0.04006 | 0.18883 | -0.33637 | -0.34668 | 0.41855 |
| Speed | 0.38437 | 0.39881 | 0.01459 | -0.03684 | -0.38532 | 0.47783 | 0.47387 | 0.10991 | -0.27966 |
| Density | -0.44780 | 0.15275 | -0.17730 | -0.28723 | -0.10494 | 0.39176 | -0.22639 | 0.59874 | 0.29662 |
| SmaCC | 0.47949 | -0.01971 | -0.02978 | 0.22827 | 0.18572 | 0.08799 | -0.60878 | 0.39804 | -0.38135 |
| Golgi intensity | 0.44387 | -0.25432 | 0.15955 | 0.21626 | 0.03223 | -0.08963 | 0.27639 | 0.39335 | 0.65260 |
| Golgi ring diameter | 0.26663 | 0.01457 | 0.33825 | -0.70830 | 0.49742 | 0.20433 | -0.02300 | -0.13399 | 0.07035 |
| Long pause CSC | -0.29580 | -0.37303 | 0.28350 | 0.41328 | 0.34322 | 0.59114 | 0.18162 | -0.02995 | -0.14506 |
| Delivery rate | 0.05254 | 0.27939 | -0.67783 | 0.13054 | 0.60742 | 0.01617 | 0.26498 | -0.00221 | 0.05693 |

**Figure S9.** Principal component analysis of cellulose biosynthesis and CSC trafficking parameters.

(**A** and **B**) Eigenvalues and Eigenvectors for principal component analysis of cellulose biosynthesis and CSC trafficking parameters, respectively. Bars in **A** indicate percent of variance that each component explains in the sample. Entries in **B** represent the correlation between each attribute (i.e., cellulose content, hypocotyl length etc.) and a principal component. Text shading emphasizes the magnitude of that correlation. (**C**) Loading plot of PC1 and PC2. Dots represent different mutations. Mutated amino acids in the conserved catalytic core motifs are highlighted with different colors. Other mutations outside of the core motifs are shown in dark grey. Proportion of variances for PC1 and PC2 are shown in parentheses. The 18 mutations could be divided into two major groups based on PC1.

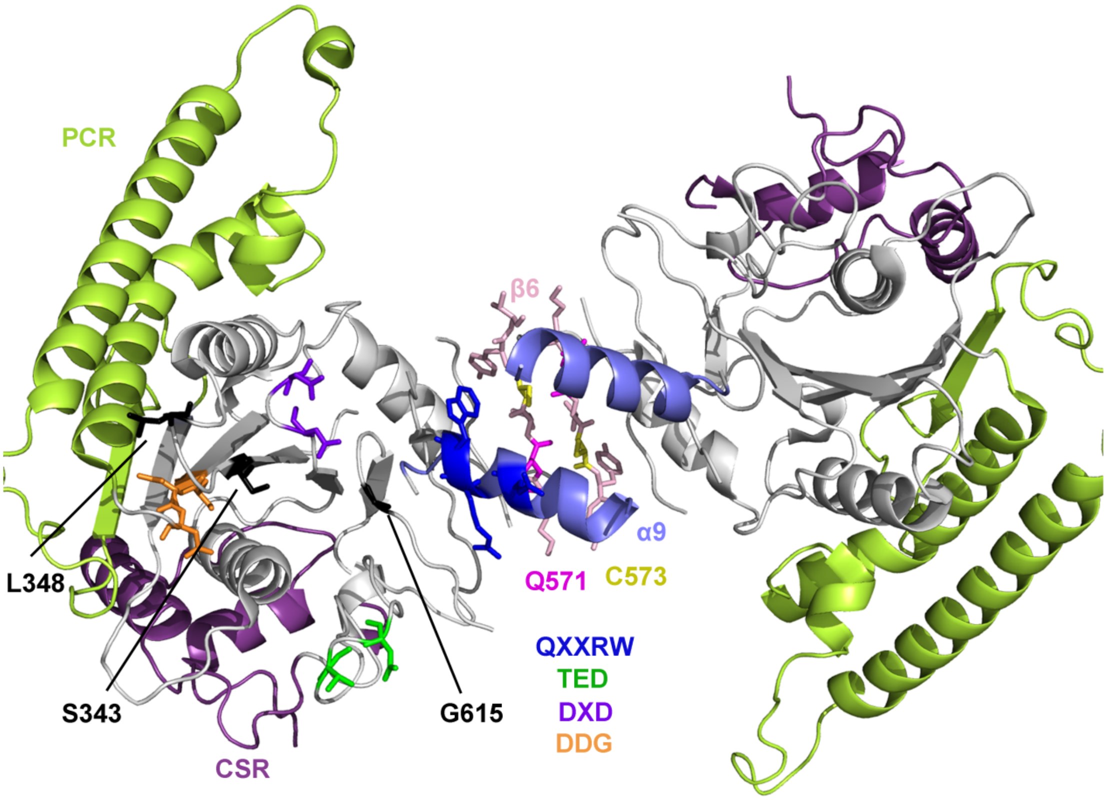


**Figure S10.** Group II homologous mutations are close to the homodimer interface formed by β-strand 6 (β6) and α-helix 9 (α9) in AtCESA3 catalytic domain structure. (Supports Figure 1)

AtCESA6 homologous mutations were mapped on the homodimer structure of AtCESA3 catalytic domain (PDB ID: 7CK1). Several amino acids are missing due to the incomplete structure of AtCESA3. Group I: CESA6-S360 (CESA3-S343), L365 (L348), S394 (S377), D395 (D378), D396 (D379), D602 (D585) (missing), D605 (D588) (missing), E929 (E909) (missing), G935 (G915) (missing). Group II: D562 (D545), D564 (D547), Q823 (Q803), R826 (R806), W827 (W807), P595 (P578) (missing), G632 (G615), T783 (T763), D785 (D765). The amino acid of AtCESA6 is given first and the homologous amino acid in AtCESA3 is shown in parentheses. Q571 and C573 are the amino acids involved in CESA3 homodimerization, as described in Qiao et al., 2021.
